# Affinity hierarchies and amphiphilic proteins underlie the co-assembly of nucleolar and heterochromatin condensates

**DOI:** 10.1101/2023.07.06.547894

**Authors:** Srivarsha Rajshekar, Omar Adame-Arana, Gaurav Bajpai, Serafin Colmenares, Kyle Lin, Samuel Safran, Gary H Karpen

## Abstract

Nucleoli are surrounded by Pericentromeric Heterochromatin (PCH), reflecting a close spatial association between the two largest biomolecular condensates in eukaryotic nuclei. Nucleoli are the sites of ribosome synthesis, while the repeat-rich PCH is essential for chromosome segregation, genome stability, and transcriptional silencing. How and why these two distinct condensates co-assemble is unclear. Here, using high-resolution live imaging of *Drosophila* embryogenesis, we find that *de novo* establishment of PCH around the nucleolus is highly dynamic, transitioning from the nuclear edge to surrounding the nucleolus. Eliminating the nucleolus by removing the ribosomal RNA genes (rDNA) resulted in increased PCH compaction and subsequent reorganization into a toroidal structure. In addition, in embryos lacking rDNA, some nucleolar proteins were redistributed into new bodies or ‘neocondensates’, including enrichment in the PCH toroidal hole. Combining these observations with physical modeling revealed that nucleolar-PCH associations can be mediated by a hierarchy of interaction strengths between PCH, nucleoli, and ‘amphiphilic’ protein(s) that have affinities for both nucleolar and PCH components. We validated this model by identifying a candidate amphiphile, a DEAD-Box RNA Helicase called Pitchoune, whose depletion or mutation of its PCH interaction motif disrupted PCH-nucleolar associations. Together, this study unveils a dynamic program for establishing nucleolar-PCH associations during animal development, demonstrates that nucleoli are required for normal PCH organization, and identifies Pitchoune as an amphiphilic molecular link required for PCH-nucleolar associations.

## Introduction

The eukaryotic nucleus is organized into different membrane-less compartments or biomolecular condensates that assemble via phase separation or similar mechanisms driven by multivalent interactions between their constituent molecules ^1,2^. An individual condensate concentrates several macromolecules, including structured and intrinsically disordered proteins and nucleic acids ^3,4^. Condensates with the same components can nucleate and grow at different cellular locations and coarsen by fusions into larger clusters, whereas those with distinct compositions do not mix ^5^. Nevertheless, in the crowded environment of the nucleus, different condensates display close, conserved associations to form higher-order complex structures ^6^. Although many studies have examined the formation and function of individual condensates, how distinct interacting condensates form and impact each other *in vivo* is less clear. This study addresses this question in the context of the two largest nuclear condensates, the nucleolus and heterochromatin.

The nucleolus is the site of ribosome synthesis with additional functions in cell cycle progression, stress response, and protein sequestration ^7^. The nucleolus assembles on chromosomal loci with transcribing ribosomal RNA genes (rDNA) and recruits specific factors involved in ribosomal RNA (rRNA) transcription, processing, and ribosome assembly to form three sub-compartments with different compositions and material properties ^8,9^. This organization is thought to facilitate the vectorial expulsion of ribosomes through and out of the nucleolus ^10,11^. In most eukaryotic nuclei, the nucleolus is surrounded by Pericentromeric Heterochromatin (PCH), a chromatin compartment composed of megabases of pericentromeric repeats, including tandemly repeated satellite DNA and transposable elements^12,13^. PCH is associated with transcriptional silencing and has essential roles in nuclear architecture, chromosome segregation, and genome stability ^14^. Under the microscope, PCH can be visualized as chromatin regions enriched for the AT-rich DNA dye DAPI, histone modifications di- and tri-methylation of histone H3 (H3K9me2/3), and the cognate epigenetic reader Heterochromatin Protein 1 (HP1) ^15^. HP1 is a multivalent protein with structured and disordered domains ^16^ that phase separates and partitions DNA and nucleosomes *in vitro* ^17,18^, and forms a liquid-like condensate nucleated by H3K9me2/3 enriched chromatin *in vivo* ^19–21^. One explanation for why the nucleolus is adjacent to PCH is that tandem repeats of rDNA are positioned next to heterochromatic satellite repeats on a subset of chromosomes ^22^. However, cytological and sequencing analyses have revealed that sequences from most chromosomes (including those lacking rDNA) make contacts with the nucleolus ^23–25^, suggesting that cis-proximity to rDNA is not necessary for PCH to organize at the nucleolar edge. The mechanisms that position PCH from all chromosomes around the nucleolus are unclear. Understanding this is important, as PCH dissociation from nucleoli in senescent cells suggests a link between PCH-nucleolar association and cellular health^26^.

In this study, we use live imaging and genetic tools in the *Drosophila melanogaster* model to uncover the dynamic patterns of *de novo* assembly of PCH from all chromosomes around the nucleolus. Removal of rDNA, and thus a functional nucleolus, caused dramatic changes in PCH assembly dynamics and redistribution of some nucleolar proteins into new bodies or ’neocondensates’. These *in vivo* phenotypes led us to develop a physical model based on a hierarchy of interaction strengths between PCH, nucleoli, and ’amphiphilic’ protein(s) able to interact with both nucleolar and PCH components. Simulations recapitulated the layered organization of nucleoli and PCH, as well as the phenotypes caused by rDNA deletion. Importantly, this model was validated by demonstrating that Pitchoune, a DEAD-box RNA-Helicase protein, acts as an amphiphilic linker responsible for PCH-nucleolar associations. We propose that disrupting affinity hierarchies between interacting condensates can redistribute their constituents to form neocondensates or other aberrant structures that may result in cellular disease phenotypes.

## Results

### Dynamic conformational changes during assembly of PCH around the nucleolus during *Drosophila* embryonic development

As in other eukaryotes, *Drosophila* nucleoli are multi-layered ^9,27^, with the outermost layer Granular Component (GC) marked by Modulo (the fly ortholog of Nucleolin) and the Dense Fibrillar Component (DFC) marked by Fibrillarin. Live imaging in late embryos (∼12-16 hr, Embryonic Stage 14-16) co-expressing fluorescently tagged transgenes of Fibrillarin or Modulo along with HP1a, shows that HP1a is positioned around the GC at the apical edge of the nucleus (**Fig. 1a, Extended Data Fig. 1a).** In the canonical ’surrounded’ conformation, HP1a does not fully engulf the nucleolus but organizes around its lateral sides, covering ∼30% of the nucleolar surface in 3D (**Supplementary Movie 1**). This surrounded conformation persists through development, as observed in gut cells in late embryos, epidermal cells in first instar larvae, and eye discs in third instar larvae (**Extended Data Fig. 1b)**.

**Fig. 1:**
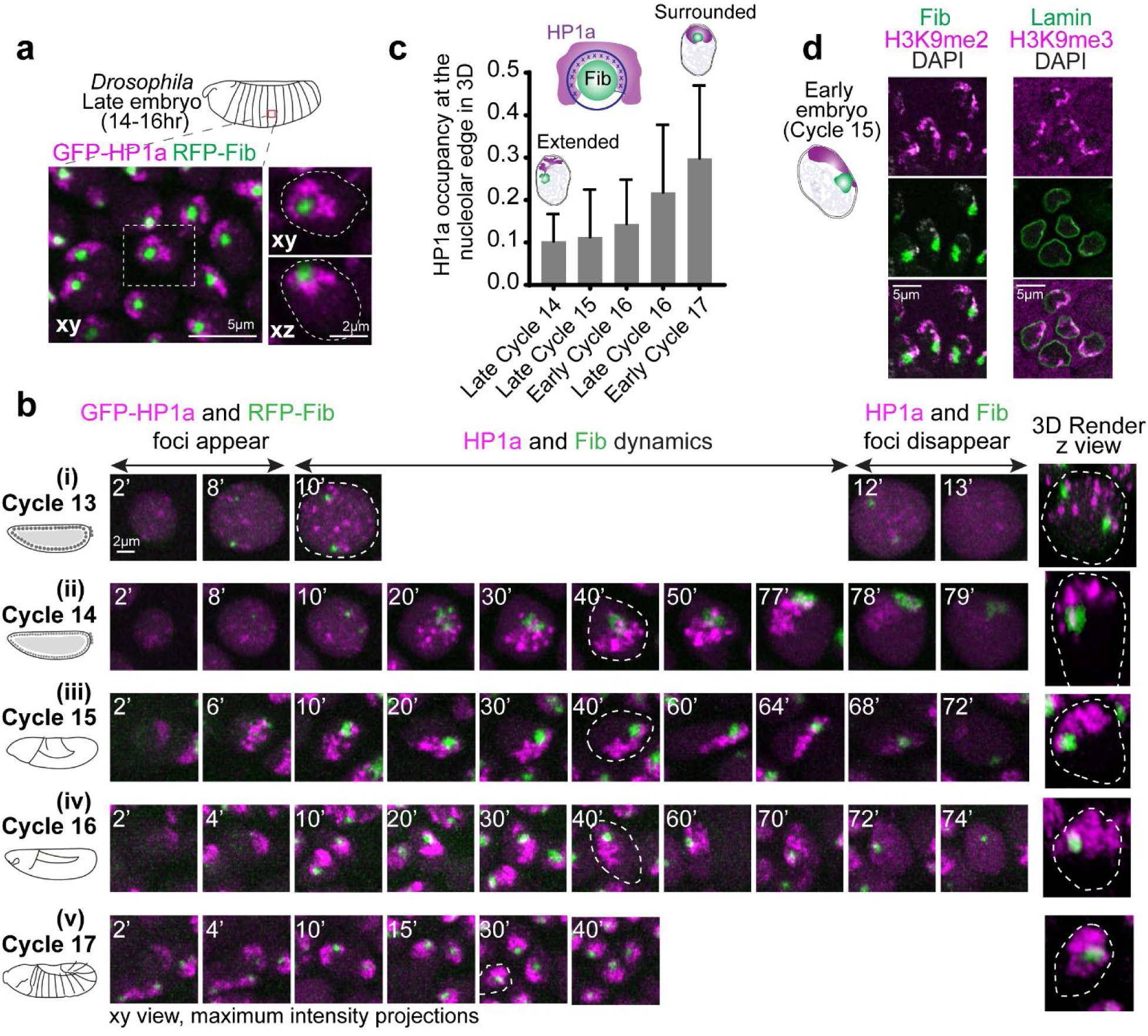
PCH is reorganized during *Drosophila* embryonic development from extended away to positioned around the nucleolus. (**a**) Maximum intensity projections showing the distribution of GFP-HP1a (magenta) and RFP-Fibrillarin (green) in live epidermal nuclei from a late-stage *Drosophila* embryo (Stage 16, ∼14-16hr). The nucleus outlined by the white dashed box is magnified on the right, presenting xy and xz views with white dashed lines indicating the nuclear boundary. (**b**) Maximum intensity projections (xy view) of individual live nuclei of GFP-HP1a (magenta) and RFP-Fibrillarin (green) localization in Cycles 13-17 of *Drosophila* embryogenesis. The numbers on the top left corner of each image indicate time (in minutes) after mitotic exit. The 3D render of the nucleus marked with white dashed lines is presented (not to scale) at the right end of the panel. (**c**) Quantification of HP1a occupancy at the nucleolar edge during specific developmental cycles. "Early Cycle" refers to nuclei between 15-30 mins into the specified interphase, "Late Cycle" refers to nuclei between 50-70 mins. Schematic above the graph illustrates the approach used to calculate the fraction of the nucleolar edge (dark blue shell) intersecting with HP1a (dark blue crosses). Bar graphs represent mean with s.d. n>50 nuclei (from 5 embryos) at each time point. (**d**) Immunofluorescence staining of nuclei from early (Cycle 15) embryos. Left: H3K9me2 (magenta) and Fibrillarin (green). Right: H3K9me3 (magenta) and Lamin (green).

To determine how PCH forms the surrounded conformation around the nucleolus, we performed high-resolution time-lapse imaging of HP1a and Fibrillarin in early *Drosophila* embryos. *Drosophila* embryos undergo 14 syncytial nuclear divisions before cellularization at the blastoderm stage, with chromatin features such as H3K9 methylation progressively established during these cycles ^28^. PCH condensates first emerge in cycle 11 ^19^ while nucleoli first emerge in cycle 13 ^29^, making cycle 13 the earliest time both condensates appear in the same nucleus. Upon entry into cycle 13 interphase, HP1a and Fibrillarin proteins are initially diffuse throughout the nucleus, then within ∼8 mins, each becomes enriched in multiple, distinct foci (**Fig. 1b(i) and Supplementary Movie 2**). Small PCH and nucleolar condensates remain separated throughout cycle 13, likely because growth is limited by the short interphase (∼15 mins) before both dissolve in mitosis ^19,29^. Cycle 14 begins like cycle 13 in that HP1a and Fibrillarin foci emerge soon after mitotic exit and are initially separated. During the longer interphase (90 min) of cycle 14, PCH and nucleoli undergo extensive growth in volume and self-fusions ^19,21,29^. However, instead of forming the canonical surrounded conformation, PCH extends away from the nucleolus while being tethered to the nucleolus at one end, hereafter referred to as the ’extended conformation’ (**Fig. 1b(ii) and Supplementary Movie 3**). The extended conformation is also observed in nuclei with two nucleoli, which appear when the two rDNA arrays in a nucleus are unpaired (**Extended Data Fig. 1c**).

Continued live imaging of HP1a and Fibrillarin in post-blastoderm, asynchronous cell divisions revealed how the extended PCH configuration dynamically transitions into the surrounded form observed in later developmental stages. PCH rapidly forms the extended configuration by lining the nuclear edge through the rest of cycle 15 (**Fig. 1b(iii) and Supplementary Movie 4**), transitions between the extended and surrounded configurations during cycle 16 (**Fig. 1b(iv) and Supplementary Movie 5**) and stably wraps around the nucleolus ∼15 min into cycle 17 interphase (**Fig. 1b(v) and Supplementary Movie 6**). HP1a occupancy around the nucleolus increases from 10% to 30% in 3D between cycles 14 and 17 (**Fig. 1c**). HP1a reorganization observed in Cycle 17 is mirrored in cultured S2 cells exiting mitosis, where HP1a transitions through an ’extended’ intermediate before stably surrounding the nucleolus (**Extended Data Fig. 1d**).

Next, we performed DNA FISH for pericentromeric repeats and rDNA to determine how PCH and nucleolar DNA sequences are reorganized in 3D during development. In the early embryo, PCH is connected to the nucleolus due to the physical proximity of rDNA repeats to pericentromeric repeats on the X and Y chromosomes in both females (XX) and males (XY) (**Extended Data Fig. 2a-b**). In the late embryo, we observed an intense rDNA signal at the nucleolar periphery, adjacent to the 359bp sequence on the X chromosome (**Extended Data Fig. 2c**). This is consistent with the process of nucleolar dominance, where the rDNA array on one X chromosome is silenced in *Drosophila melanogaster* ^30,31^, and with the repositioning of silent rDNA outside the nucleolus ^32^. The 1.686 repeat sequences, located on chromosomes lacking rDNA (chromosomes 2 and 3), are positioned away from the nucleolus during cycle 15 (Stage 8), forming the ’extended conformation’, but relocate to the nucleolar edge in the late embryo (**Extended Data Fig. 2d-e**). Immunostaining early embryos for H3K9me2/3, Lamin, and Fibrillarin reveals that the PCH lines the nuclear lamina in the ’extended conformation’ (**Fig. 1d**).

Together, these experiments detail the reorganization of the two largest biomolecular condensates during *Drosophila* embryonic development at high spatial and temporal resolution. We observe that PCH and nucleolar condensates undergo independent nucleation, growth, and fusion, displaying cycle-specific differences while dynamically transitioning from the extended (predominantly nuclear periphery associated) to the surrounded configurations (nucleoli associated) (summarized in **Extended Data Fig. 2f**).

### Embryos lacking nucleoli display increased PCH compaction and neocondensate formation

The specific patterns of PCH reorganization around the nucleolus during embryonic development prompted us to ask if the nucleolus impacts PCH assembly dynamics. We imaged RFP-HP1a and GFP-Fib in embryos that lack any rDNA repeats (designated hereafter as -rDNA) due to a rearranged X chromosome (C(1)DX/0). No functional nucleoli are formed in -rDNA embryos ^29^, but they develop through early embryogenesis due to maternal deposition of ribosomes. Fibrillarin forms spherical structures (as previously reported by Falahati et al. 2016 ^29^), which are located at a significantly longer distance from the HP1a/PCH condensate compared to their distance in wildtype (+rDNA) controls (**Fig. 2a-b**; +rDNA mean=1.78*μ*m, -rDNA mean=3.69*μ*m, p<0.0001). This result suggests that Fibrillarin and HP1a proteins do not interact directly. Instead, when factors responsible for nucleolus formation (rDNA/rRNA) are not present, Fibrillarin self-associations ^33^ and/or secondary affinities with other structures or molecules are sufficient to cause the formation of new enrichments (hereafter termed ’neocondensates’) not found in wildtype cells. The ’extended’ HP1a conformation typically observed in cycles 14 and 15 is replaced by a collapsed, rounded structure at the apical end of -rDNA nuclei (**Fig. 2a, c**). The aspect ratio (major axis/minor axis) of the HP1a domain is significantly reduced (**Fig. 2d**; +rDNA mean=1.96, -rDNA mean=1.40, p<0.0001). Additionally, the distance between pericentromeric repeats 1.686 (on Chr 2 and 3) and AAGAG (satellite repeats on all chromosomes) is decreased in -rDNA nuclei compared to +rDNA controls (**Fig. 2e-f**; +rDNA mean=0.74*μ*m, -rDNA mean=0.32*μ*m, p<0.0001). Thus, PCH loses the extended configuration and shows increased compaction in the absence of rDNA/nucleoli, where compaction is defined by the reduction in 3D space between PCH elements.

**Fig. 2:**
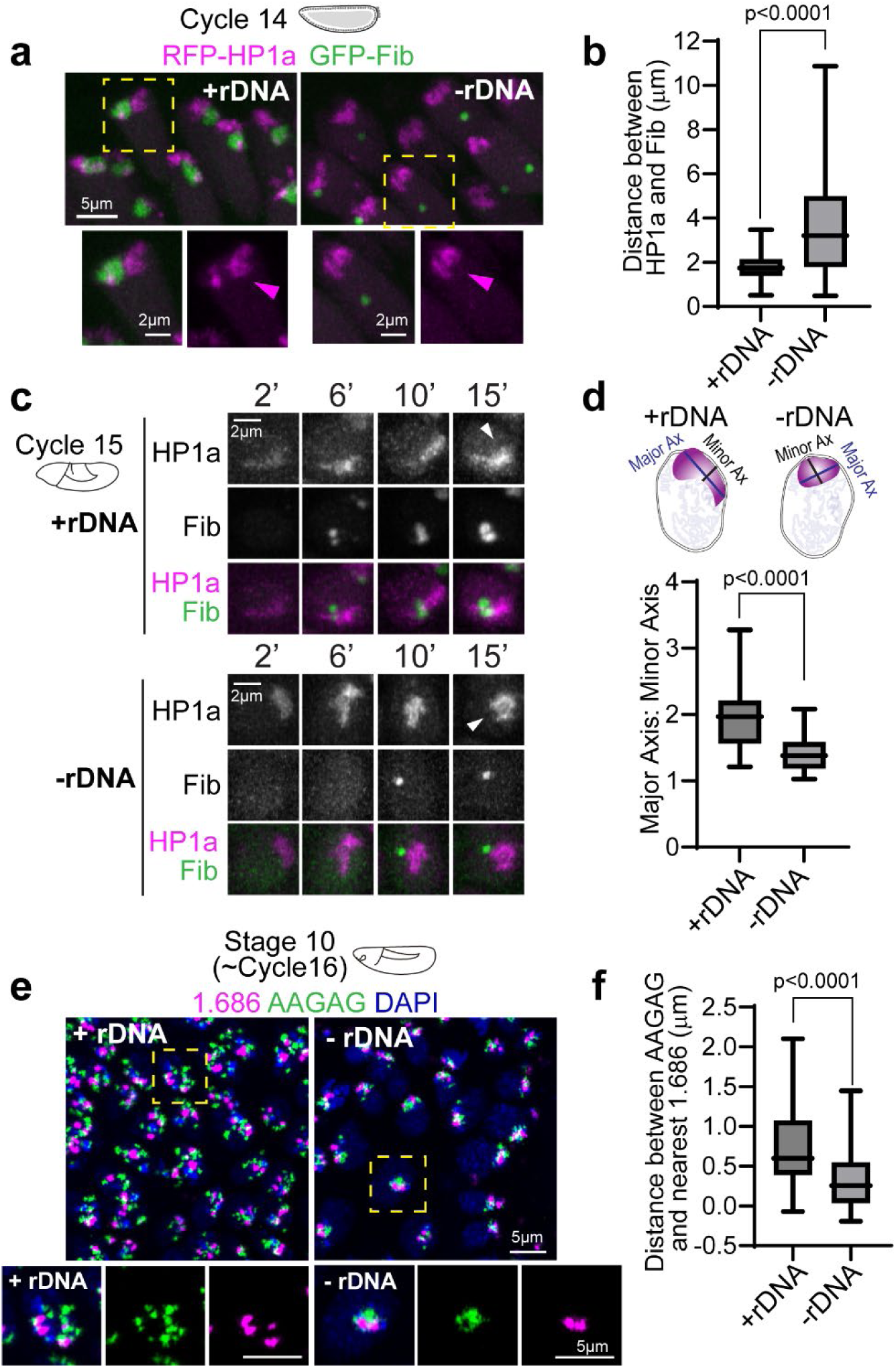
Increased PCH compaction in embryos lacking rDNA. (**a**) Maximum intensity projections of live nuclei from wildtype embryos (+rDNA) and mutant embryos lacking rDNA (-rDNA) at Cycle 14, expressing RFP-HP1a (magenta) and GFP-Fibrillarin (green). The dashed yellow box highlights the nucleus enlarged in the panel below. (**b**) Distance between the centers of geometry of HP1a and Fibrillarin in +rDNA and -rDNA embryos. n>200 nuclei (from 5 embryos) for each genotype. (**c**) Maximum intensity projections from live imaging of nuclei from +rDNA and -rDNA embryos showing the reassembly of RFP-HP1a (magenta) and GFP-Fibrillarin (green) at the indicated minutes after the start of Cycle 15. White arrowheads point to HP1a organization 15 minutes after the start of Cycle 15 in +rDNA and -rDNA embryos. (**d**) Quantification of the aspect ratio (major axis over minor axis) of HP1a segments 15 minutes after the start of Cycle 15 in +rDNA and -rDNA embryos. n>30 nuclei in each genotype. (**e**) Maximum intensity projections of FISH (Fluorescence in Situ Hybridization) of satellite repeats 1.686 (magenta) and AAGAG (green) in DAPI (blue)-stained nuclei in +rDNA and -rDNA Stage 10 embryos. The dashed boxes mark the nuclei enlarged on the right. (**f**) Quantification of the distance between AAGAG and its nearest 1.686 locus. n>40 pairs of loci in each genotype. Box plots extend from 25^th^ to 75^th^ percentile. Error bars: Min to Max.

To our surprise, PCH transitioned to a toroidal (donut-like) structure in all imaged cells in late-stage -rDNA *Drosophila* embryos, with a core devoid of HP1a (’PCH void’) (**Fig. 3a**). We visualized this transition at increased spatial resolution in the large nuclei of the amnioserosa, which forms a monolayer on the dorsal surface of the embryo during gastrulation ^34^ (**Fig. 3b-c and Supplementary Movie 7**). In addition to the absence of HP1a, the PCH void also lacked the major nucleolar proteins Fibrillarin (**Fig. 3c**), which formed a separate neocondensate, and Modulo, which dispersed in the nuclear space (**Extended Data Fig. 3a**). The PCH void did not contain DNA (DAPI staining), histones marked with H3K9me2 (IF) (**Extended Data Fig. 3c**), or any significant accumulation of RNAs (Propidium Iodide staining) (**Extended Data Fig. 3d**). However, treatment with 488 NHS Ester, a pan-protein label ^35^, revealed that the PCH void was enriched for proteins (**Fig. 3d**).

**Fig. 3:**
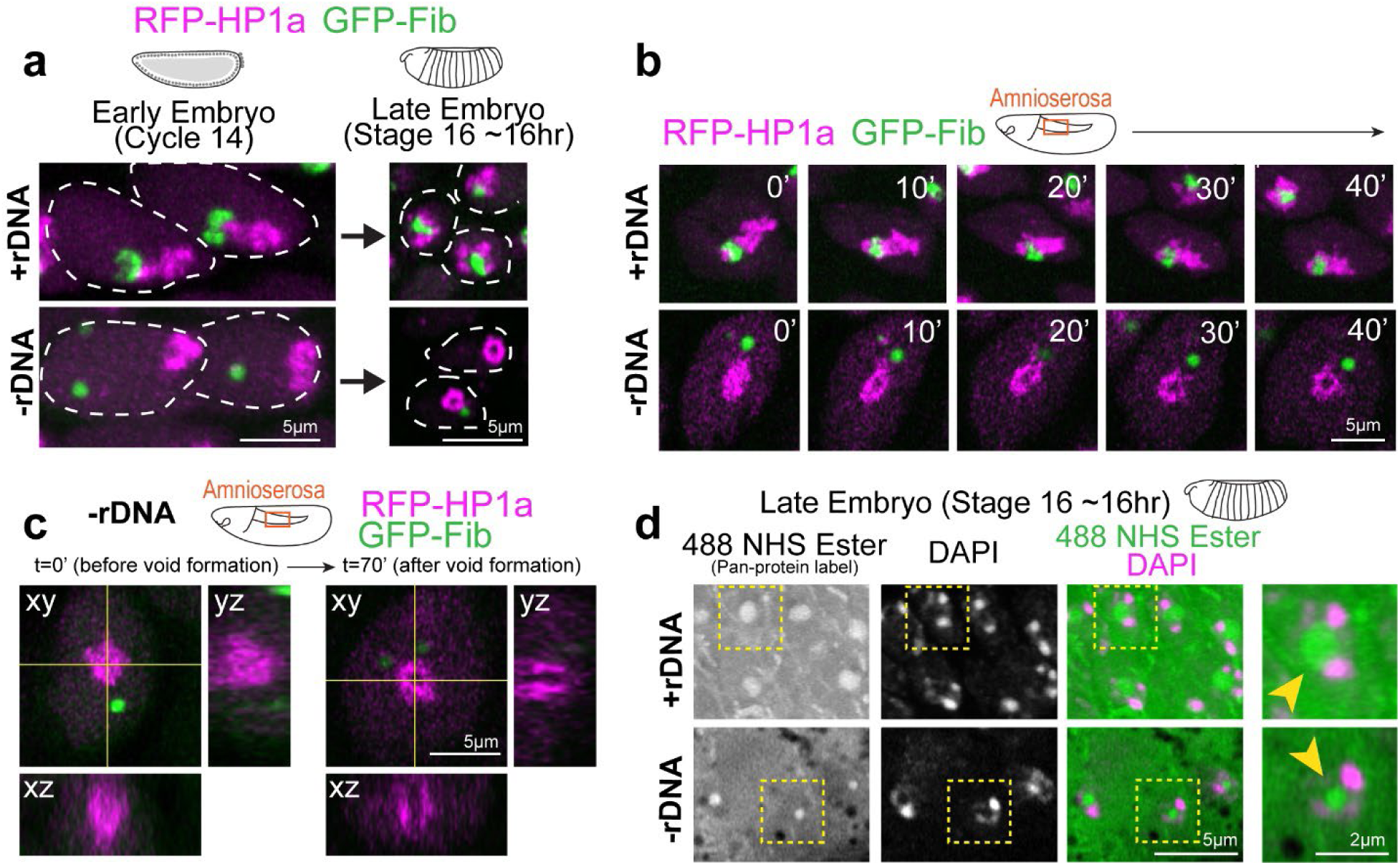
A protein-filled core reshapes the PCH condensate to a toroid-like structure in -rDNA developing embryos. (**a**) Representative stills of live +rDNA and -rDNA nuclei at Cycle 14 and in Stage 16 (late stage) embryos showing RFP-HP1a (magenta) and GFP-Fibrillarin (green). (**b**) Time-lapse stills (single slices) of a +rDNA and -rDNA amnioserosa nucleus with HP1a (magenta) and Fibrillarin (green). T=0 mins was set to capture the time window where PCH transitions from a compacted to a toroidal structure in -rDNA amnioserosa nuclei. (**c**) Orthogonal projections along yellow intersecting lines in an amnioserosa nucleus from a -rDNA embryo before and after the formation of the ‘PCH void’. (**d**) Late embryos (Stage 16) with the +rDNA or -rDNA genotype co-stained with a pan-protein label, 488 NHS Ester (green), and DAPI (magenta) show that the ‘PCH void’ in -rDNA embryos is enriched for proteins (yellow arrowhead). Nuclei marked with the yellow dashed box are enlarged on the right.

We conclude that PCH initially displays an atypical, compacted morphology in embryos lacking rDNA and nucleoli. As development proceeds, the PCH morphs into an abnormal toroidal structure whose central core (the PCH void) lacks HP1a, nucleolar proteins, chromatin, and RNA but is filled with protein(s) that may represent another neocondensate. Together, these results reveal that nucleoli are required to organize PCH in the 3D nuclear space by preventing PCH hyper-compaction and that disrupting interactions within or between condensates can create new nuclear structures.

### Coarse-grained modeling recapitulates *in vivo* PCH-nucleolar organization phenotypes and highlights a potential role for amphiphilic proteins in mediating their association

Since the nucleolus and PCH assemble via phase separation or similar mechanisms ^8,19,21,36^, we drew inspiration from the physical theory of three-phase wetting to better understand PCH-nucleolar organization in the nucleoplasm. This theory states that an equilibrium between the interfacial tensions (i.e., the energetic cost of forming an interface between two phases) determines the spatial configurations of the different phases ^37^ (**Fig. 4a**, fully engulfing, partial engulfing, or individual separated phases). These interfacial tensions are dictated by the relative interaction strengths between the components of the different phases. To describe the different three-phase spatial configurations we introduced the spreading coefficient of a phase *i*, *S_i_*,

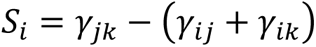

**Fig. 4:**
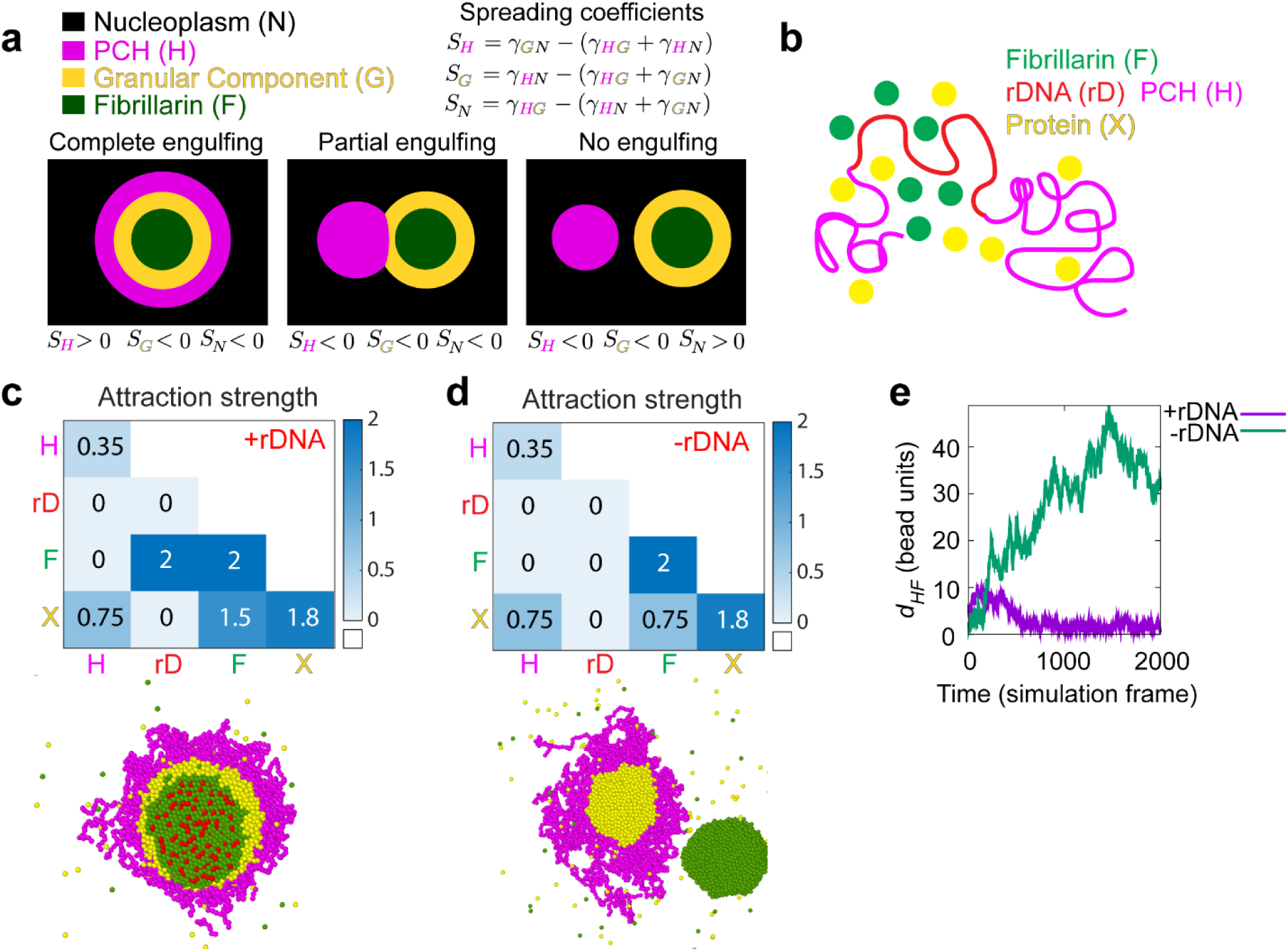
A hierarchy of interaction strengths between PCH, nucleoli and ‘amphiphilic’ protein(s) recapitulates their +rDNA and -rDNA *in vivo* organization. (**a**) The schematic shows the spatial organization of three different phases. The interfacial tensions, *γ_ij_*, between phases *i* and *j* (where *i* and *j* can be nucleoplasm (N) in black, PCH (H) in magenta, or the granular component (G) in yellow) are used to define the spreading coefficients, *Si*, shown in the top right corner. Given the assumption that *γ_GN_>γ_HN_*, the three possible combinations of spreading coefficients determine the configuration of the three phases, ranging from complete engulfing to partial engulfing to no engulfing. Since the Fibrillarin-rich phase (F) is always within a functional nucleolus, its spreading coefficients are not included in this analysis. (**b**) Course-grained modelling of nucleolar-PCH assembly with four minimal components: PCH (H) as a self-interacting polymer (magenta) rDNA (rD) as polymeric block embedded within PCH (red), Fibrillarin (F) as a representative of a self-associating nucleolar protein (green), and a self-associating protein ‘X’ (yellow) representing those enriched in the PCH void in -rDNA nuclei. (**c** and **d**) The matrices indicate the strengths of interaction (blue gradient, units: k_B_T) between Fibrillarin (F) (green), rDNA (rD) (red), PCH (H) (magenta), and protein (X) (yellow), which has dual affinities for both H and F. The indicated affinity hierarchies result in simulated outcomes that recapitulate their +rDNA and -rDNA organization observed *in vivo*. (**e**) Distance (*d_HF_*) between the centers of mass of PCH (H) and Fibrillarin (F) clusters in +rDNA and -rDNA simulations.

where the letters *i, j*, and *k*, represent three different phases, *γ*_*jk*_ is the interfacial tension between phases *j* and *k γ*_*ij*_ is the interfacial tension between phases *i* and *j*, and *γ*_*ik*_ is the interfacial tension between phases *i* and *k*. For every three-phase combination, there are three spreading coefficients, and depending on whether such spreading coefficients are positive or negative, different spatial configurations are attained ^37^. We exemplify this using PCH (H), the outermost nucleolar layer GC (G), and the nucleoplasm (N) (**Fig. 4a**).

To connect the theory of wetting with our *in vivo* results, we used a minimal coarse-grained model that incorporates the molecular interactions and biophysical parameters that could mediate nucleolar-PCH spatial organization with four minimal components (**Fig. 4b**). (i) PCH (H) as a long, self-attracting polymer. (ii) Fibrillarin (F), which represents a self-associating nucleolar protein. (iii) rDNA (rD) as a polymeric block within the chromatin fiber that experiences good solvent conditions and is flanked by two PCH blocks. Although we do not explicitly account for the presence of rRNA, we use rDNA as a proxy for the nucleating site for Fibrillarin condensates. Finally, (iv) Protein (X), as a self-associating molecule with an affinity for PCH. These properties were attributed to protein X to account for the spherical protein-rich compartment that consistently formed within the PCH-void in -rDNA nuclei (**Fig. 3d**). We then considered what additional properties Protein X could possess to form the observed *in vivo* organization of the nucleolus and PCH.

To recapitulate the layered PCH-nucleolar organization, we reasoned the self-associations of Fibrillarin must be stronger than those of protein X, which in turn must be stronger than the self-interactions of PCH. These self-interaction choices result in the following hierarchy of interfacial tensions *γ*_*FN*_ > *γ*_*XN*_> *γ*_*HN*_, where *γγ*_*iN*_ is the interfacial tension between a phase enriched in component *i = F, X, H* and the nucleoplasm (*N*). Additionally, we note that for a stable association between PCH and the Fibrillarin-rich phase, Protein X must also have an affinity for Fibrillarin (or more generally for nucleolar components), which *in vivo* might be mediated by the presence of rRNA. Therefore, we define Protein X as an ’amphiphilic protein’, due to its dual affinity for the PCH and nucleolar phases, as defined previously for synthetic co-condensates ^38^. Altogether, these considerations led us to define a hierarchy of interaction strengths (rD-F ≥ F-F > X-X > F-X > X-H > H-H) as an initial set of parameters listed in the 4×4 interaction matrix (**Fig. 4c**). Simulating this affinity hierarchy recapitulated the canonical PCH-nucleolar organization observed in +rDNA animals *in vivo* (**Fig. 4c and Supplementary Movie 8**). Various scenarios incompatible with the experimental observations of the layered organization of the nucleolus and PCH arise by deviating from the hierarchy of self-interaction strengths in the simulations. For example, instead of having a complete engulfing of the nucleolus by PCH, there are partial engulfing or no engulfing scenarios when the interaction between the amphiphilic proteins is larger than that of Fibrillarin (**Extended Data Fig. 4d**). Importantly, our choice of interaction parameters is not unique as long as the interfacial tensions fulfill *γ*_*FN*_ > *γ*_*XN*_ > *γ*_*HN*_, we obtain a complete engulfing of the nucleolus by PCH. A detailed rationale for the choice of parameters is described in the Methods section.

Next, we modeled the consequences of removing rDNA by setting the rDNA-Fibrillarin interactions to zero, thus eliminating its ability to recruit nucleolar components such as Fibrillarin and X. Reducing the attraction between Fibrillarin and the amphiphilic protein X, which increases the interfacial tension between the Fibrillarin-rich and the amphiphilic-rich phases, *γ*_*FX*_, resulted in the formation of a protein X neocondensate within the PCH void and a spatially separated Fibrillarin neocondensate (**Fig. 4d-e**), as observed in -rDNA embryos *in vivo* (Fig. 2 and 3). These phenotypes are observed in simulations whenever the protein X – PCH interaction strength is greater than or equal to the X – Fibrillarin interactions (i.e., X-H ≥ F-X) (**Extended Data Fig. 4e-f**). In the model, the nucleation of the new phase of amphiphilic protein X occurs within the PCH to reduce the cost of creating an interface of protein X with the solvent. Instead, being surrounded by PCH can reduce this interfacial energy, as expected from a wetting picture of simple liquids ^37^. We conclude that incorporating affinity hierarchies and an amphiphilic protein X into a minimal model recapitulates the observed *in vivo* organization of PCH and nucleolar condensates in both +rDNA and -rDNA conditions. This led us to predict that in WT conditions, protein X would be a self-associating protein with dual affinities for the nucleolus and PCH, albeit with a weaker affinity for PCH components (F-X > X-H).

### The DEAD-box RNA Helicase Pitchoune is a granular compartment protein that forms a neocondensate within PCH in -rDNA embryos

A candidate for an amphiphilic protein that can interact with both PCH and the nucleolus emerged from Falahati et al., 2017 ^36^. They reported that the DEAD-box RNA helicase Pitchoune (Pit), typically a granular component (GC) nucleolar protein ^39^, mislocalized to the apical part of the nucleus in cycle 14 -rDNA embryos, similar to the compacted PCH domain observed in our studies (Fig. 2a). Pitchoune, the *Drosophila* ortholog of DDX18, is required for larval development ^39^, and possesses an N-terminal intrinsically disordered region (IDR), a central helicase core, and a disordered C-terminal domain (**Fig. 5a**). We identified two tandem "PxVxL" HP1a-interacting motifs ^40,41^ in the C-terminal tail of Pitchoune with PVVDL being conserved across eukaryotes along with LKVGA being conserved among Drosophilids (**Fig. 5a and Extended Data Fig. 5a**). Additionally, Pitchoune belongs to the DDX family of proteins, other members of which phase separate and modulate condensate behavior ^42^. Together, these clues led us to hypothesize that Pitchoune could be a candidate amphiphilic protein that can 1) form condensates and 2) have a dual affinity for both the nucleolus and PCH.

**Fig. 5:**
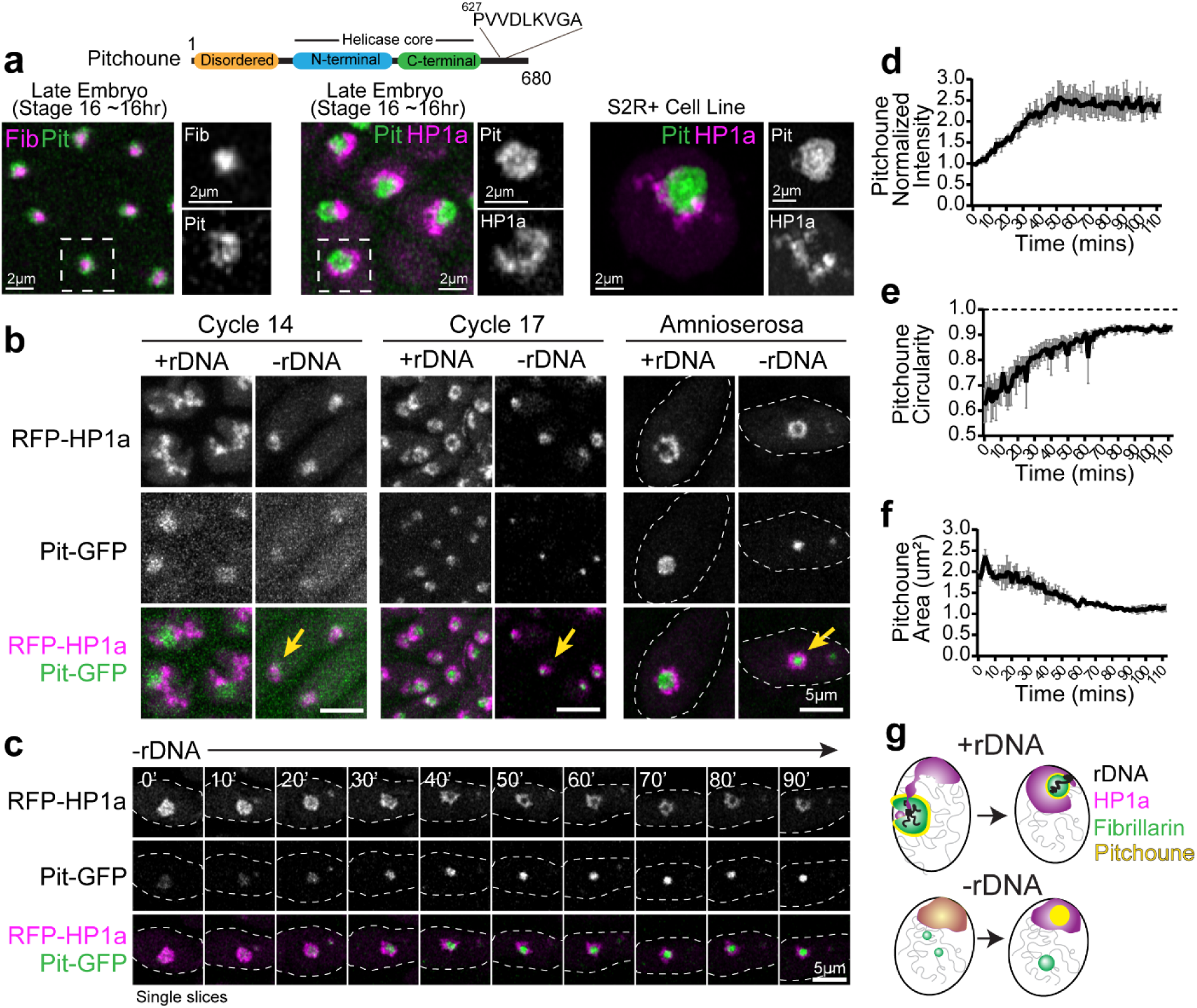
The RNA Helicase Pitchoune is enriched in the PCH void in -rDNA embryos. (**a**) Protein Subdomains in Pitchoune (Pit) with an N-terminus disordered domain, central helicase core and C-terminus with a conserved HP1a interacting motif. Left to Right: Panels showing the localization of RFP-Fib (magenta) and Pit-GFP (green) in Stage 16 embryo, RFP-HP1a (magenta) and Pit-GFP (green) in Stage 16 embryo, and Scarlet-I-HP1a (magenta) and Pit-mYFP (green) transiently transfected in S2R+ *Drosophila* cells. (**b**) Maximum intensity projections of Cycle 14, Cycle 17 and amnioserosa nuclei from +rDNA and -rDNA embryos showing RFP-HP1a (magenta) and Pit-GFP (green). Yellow arrows in -rDNA in Cycle 17 show the mixing of Pit and HP1a, while yellow arrows in -rDNA in cycle 17 and amnioserosa show the formation of the Pit neocondensate in the PCH void. (**c**) Time-lapse stills (single slices) of a -rDNA amnioserosa nuclei with RFP-HP1a (magenta) and Pit-GFP (green). (**d**) Mean intensity, (**e**) Circularity, and (**f**) Area of projections of Pitchoune in -rDNA embryos in amnioserosa nuclei. Mean of 35-60 nuclei at each time point in n=3 embryos. Error Bars: s.e.m. (**g**) Schematic summarizing nuclear organization phenotypes observed in +rDNA and -rDNA embryos.

Consistent with its role as a GC nucleolar protein ^43^, Pitchoune is enriched in the outermost nucleolar compartment in *Drosophila* embryos and S2R+ cells (**Fig. 5a**). Next, to investigate whether Pitchoune is enriched in the PCH void in -rDNA embryos, we performed live imaging in +rDNA and -rDNA embryos co-expressing Pit-GFP and RFP-HP1a transgenes. In cycle 14 +rDNA nuclei, Pitchoune localizes to the nucleolus while HP1a is in the extended configuration. In contrast, a faint Pitchoune signal co-localizes with the more compact HP1a domain in -rDNA nuclei (**Fig. 5b**). In cycle 17 epidermal nuclei and amnioserosa nuclei, HP1a surrounds Pitchoune in +rDNA embryos, whereas in -rDNA embryos, high-intensity Pitchoune puncta appear within each compacted HP1a condensate (**Fig. 5b**). Time-lapse imaging of amnioserosa nuclei in -rDNA embryos revealed that Pitchoune is initially faintly mixed but gradually separates from HP1a, with the intensity of Pitchoune increasing ∼2.5-fold over an hour (**Fig. 5c-d and Supplementary Movie 9**). Furthermore, the circularity of Pitchoune increases and approaches 1 (in projections) (**Fig. 5e**), and the area of Pitchoune decreases over time (**Fig. 5f**). These results suggest that Pitchoune demixes from HP1a and forms a spherical neo-condensate surrounded by PCH, minimizing interfacial energy by avoiding the creation of a larger energy interface between Pitchoune and the nucleoplasm (**Fig. 5g**).

### Pitchoune and its HP1a-interacting motif are required for PCH-nucleolar associations

Supported by our model and *in vivo* imaging, our findings indicate that Pitchoune is a candidate amphiphile between PCH and nucleoli as a granular compartment nucleolar protein with a lower affinity for PCH/HP1a. We next investigated the consequences of removing Pitchoune on PCH-nucleolar associations. If Pitchoune regulates these associations, we predicted that its depletion would disrupt the surrounding configuration. We first tested this by decreasing the concentration of the amphiphilic protein in the physical model while keeping all other parameters the same as in Fig. 4c. The simulations revealed a progressive detachment of PCH from the nucleolus as Pitchoune levels decreased (**Fig. 6a**). Note that in both simulations and *in vivo* experiments, even with the loss of Pitchoune, nucleoli and PCH are expected to maintain a singular attachment at the PCH-embedded rDNA locus. This is in contrast with the separation of Fibrillarin and PCH observed in embryos lacking rDNA (**Fig. 2a, 4c**).

**Fig. 6:**
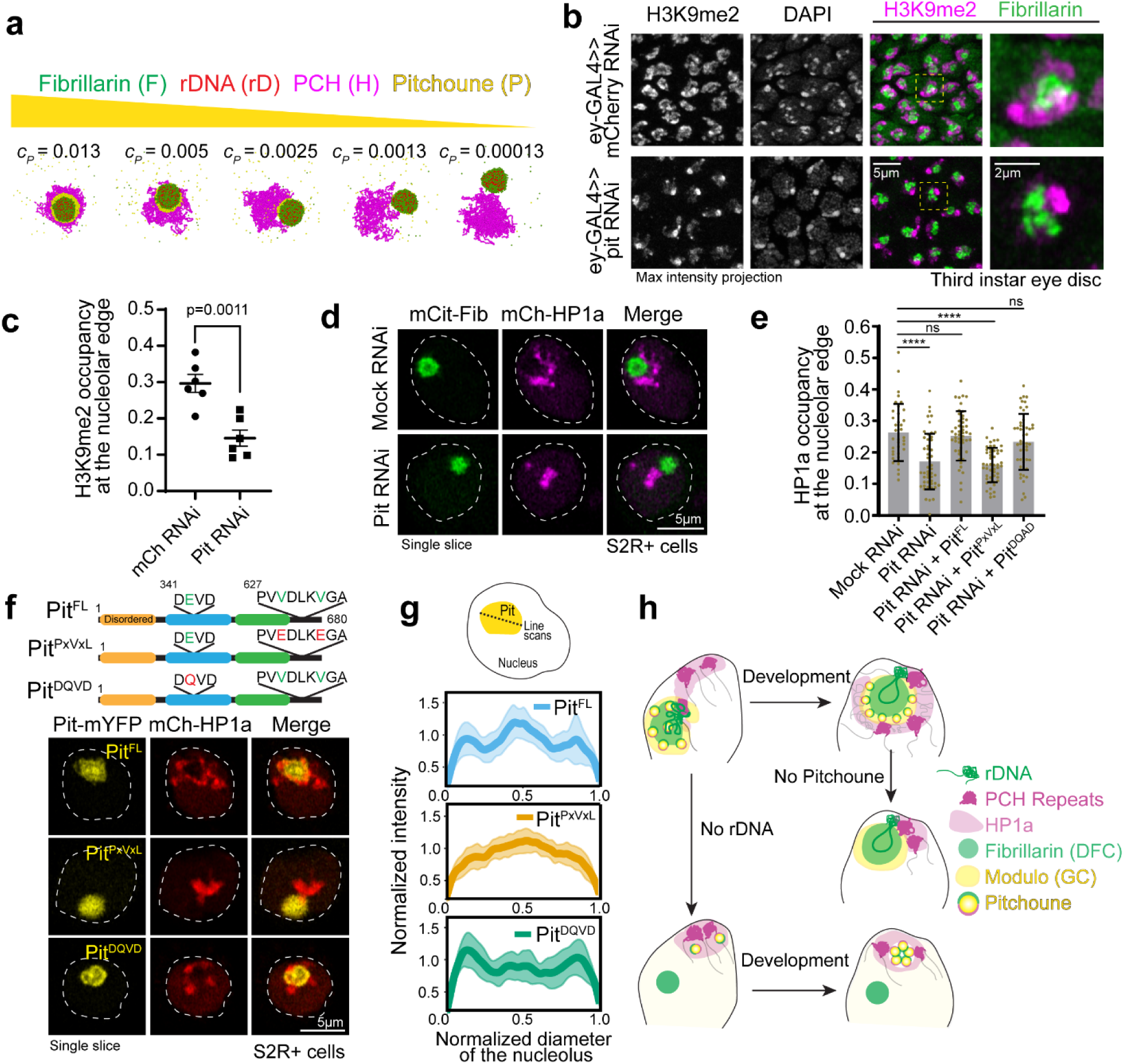
Knockdown of Pitchoune or mutations in its HP1a-interaction motif disrupt PCH-nucleolar associations. (**a**) Simulation endpoint snapshots demonstrate how decreasing the concentration (c_P_) of Pitchoune (P) (yellow) reduces the association of PCH (H) (magenta) around the Fibrillarin condensate (F) (green). (**b**) Immunofluorescence of H3K9me2 (magenta), Fibrillarin (green) and DAPI in nuclei from dissected eye-discs in third instar larvae from UAS-pit RNAi and UAS-mCherry RNAi (control) driven by the eyeless-GAL4 driver. (**c**) Quantification of the fraction of the nucleolar edge occupied by H3K9me2. Each data point indicates the mean value from one animal. n=6 animals with over 500 nuclei in total for each experimental group. Error bars: s.e.m. (**d**) Nuclei expressing Citrine-Fib and mCherry-HP1a transiently transfected after Pitchoune knockdown show decreased association of HP1a from the nucleolus. (**e**) Fraction of the nucleolar edge occupied by HP1a following Pit knockdown and rescue with Full length Pit (Pit^FL^), Pit^PxVxL^ and Pit^DQVD^. n=38-50 nuclei for each experimental group. Bar graphs show the mean ± s.d. (**f**) Schematic representation of Pitchoune mutations introduced: Pit^PxVxL^ contains mutations of the central valine (V) to glutamic acid (E) in the conserved PxVxL motifs. The putative helicase mutant is denoted as Pit^DQVD^. Representative nuclei transfected with Pit^FL^, Pit^PxVxL^, and Pit^DQVD^ following RNAi-mediated knockdown of endogenous Pitchoune. (**g**) Line scans show Pitchoune distribution in the nucleolus of cells transfected with Pit^FL^ (n=25), Pit^PxVxL^ (n=37) and Pit^DQVD^ (n=40) after Pit RNAi. Intensities are normalized to the average value for each profile, and lengths are normalized to the diameter of the corresponding nucleolus. Solid line: mean, shaded error region: s.d. (**h**) Model for the organization of PCH (repeats and HP1a) around the nucleolus (rDNA, Fibrillarin and Modulo) via an amphiphilic linker Pitchoune during *Drosophila* development.

Next, we depleted Pitchoune using RNAi in the female germline (using a maternal GAL4 driver) to assess the impact of Pitchoune knockdown in *Drosophila* embryos. However, ovary development was halted, and no eggs were laid following Pitchoune knockdown in the female germline (**Extended Data Fig. 6a-b**), preventing further assessment of the effects of Pitchoune knockdown in early embryos. These phenotypes align with the growth defects observed in *pitchoune* homozygous mutants that die as small first instar larvae ^39^, likely surviving early development due to maternal deposits of Pitchoune. Therefore, we used eyeless-GAL4 to knock down Pitchoune in the eye-antennal discs of third-instar larvae. Pitchoune knockdown severely disrupted the normal development of eye-antennal discs (**Extended Data Fig. 6c**). Most importantly, viable cells displayed a 50% reduction in the fraction of the nucleolar edge occupied by H3K9me2 after Pitchoune knockdown (control RNAi mean = 0.296, Pitchoune RNAi mean = 0.146, p-value = 0.0011) (**Fig. 6b-c**).

We also knocked down Pitchoune in S2R+ *Drosophila* cultured cells using RNAi, achieving ∼60% reduction in Pitchoune transcripts (**Extended Data Fig. 6d**). As observed in eye discs, Pit knockdown decreased HP1a occupancy at the nucleolar edge and disrupted its organization relative to the nucleolus, with ∼50% reduction in HP1a occupancy at the nucleolar edge, and a decrease in the percentage of nuclei displaying the surrounded configuration (Mock RNAi: 87%, Pit RNAi: 27%) (**Fig. 6d-e, Extended Data Fig. 6e-f**). These phenotypes were rescued by reintroducing full-length Pitchoune (**Fig. 6e-f, Extended Data Fig. 6f**). We conclude that Pitchoune is important to establish or maintain the surrounded configuration.

To determine the molecular basis of Pitchoune’s ’amphiphilic’ nature and test whether it is directly responsible for PCH-nucleolar associations, we investigated the roles of Pitchoune’s conserved motifs in mediating these interactions. We disrupted the putative HP1a-interacting PxVxL motifs by mutating the two central valines (V) to glutamic acid (E) (Pit^PxVxL^). The central V binds in a hydrophobic pocket at the interface of HP1 dimers, and mutating the hydrophobic V to the charged E residue disrupts their HP1 interaction ^40^. We also generated a construct mutating the conserved DEVD motif in the Helicase domain to DQVD (Pit^DQVD^) to block its Helicase activity ^39,44^ (**Fig. 6f**). Defects in HP1a organization relative to the nucleolus caused by Pit knockdown in S2R+ cells were rescued by transfection with full-length Pitchoune (Pit^FL^) and Pit^DQVD^ but not Pit^PxVxL^ (**Fig. 6e-f)**. When expressing Pit^FL^ and Pit^DQVD^, 73% and 62% of nuclei, respectively, exhibited the ’surrounded’ PCH-nucleolar organization after Pit knockdown, compared to only 30% of nuclei expressing Pit^PxVxL^ (**Extended Data Fig. 6f**). Additionally, Pit^PxVxL^ showed uniform distribution in the nucleolus, unlike Pit^FL^ and Pit^DQVD^ which were enriched at the nucleolar edges (**Fig. 6f, g**). We conclude that the PxVxL motif regulates Pitchoune’s sub-nucleolar distribution by promoting its enrichment at the nucleolar-PCH interface. Together, these experiments reveal the molecular basis of Pitchoune’s amphiphilic function, showing that it promotes PCH-nucleolar associations through interactions with HP1a via its PxVxL motif.

## Discussion

This study presents four main findings on the organization of PCH and nucleoli (**Fig. 6h**): (**1**) During early embryonic development, PCH-nucleolar associations are highly dynamic, transitioning from extended (nuclear edge-associated) to surrounded (nucleolar edge-associated) configurations (**Fig. 1**). (**2**) The nucleolus organizes PCH components in the 3D nuclear space, preventing PCH hyper-compaction (**Fig. 2**). (**3**) A hierarchy of interaction strengths between nucleolar and PCH components, including amphiphilic proteins with affinities for both, recapitulates the spatial organization seen in cells with and without nucleoli, and disrupting these hierarchies generates neo-condensates (**Fig. 3-5**). (**4**) Pitchoune and its C-terminal HP1a binding PxVxL motif are required for normal PCH-nucleolar associations (**Fig. 6**).

Sequence-based approaches have revealed that nuclear organization transitions from a naïve state to specific higher-order patterns during embryonic development ^45^. However, these methods typically exclude the highly repetitive sequences that comprise most of the PCH and nucleoli, limiting our understanding of how PCH-nucleoli organization is established during development. Our study addresses this gap by visualizing PCH and nucleolar dynamics during *Drosophila* embryonic development in single nuclei at minute-scale resolution. Initially, PCH from different chromosomes undergo liquid-like fusions ^19,21^ to form a contiguous condensate that wraps around the nucleolus, explaining how PCH from chromosomes without rRNA genes is also positioned at the nucleolar edge. After fusing into a contiguous condensate, PCH remains dynamic, transitioning from an ’extended’ configuration in cycle 14 to a stable ’surrounded’ state around the nucleolus in cycle 17. Transitions in heterochromatin organization reminiscent of the ‘extended’ intermediate have been observed during early embryonic development in *C. elegans* ^46^ and mice ^47^, suggesting similarities across species.

In its ’extended’ intermediate state, PCH associates closely with the nuclear edge before repositioning around the nucleolus. This finding is intriguing because sequencing-based approaches have shown overlap between Nucleolus Associated Domains (NADs) and Lamin-Associated Domains (LADs) ^24,48^. The molecular mechanisms behind this repositioning are unknown, but we speculate that PCH’s affinity for the nuclear periphery decreases while its interaction strength with the nucleolus increases between cycles 14 and 17, with Pitchoune playing a crucial role. Post-translational modifications or increased concentrations of Pitchoune in cycle 17 might enhance its affinity for HP1a, correlating temporally with the stable surrounding of the nucleolus by HP1a in +rDNA and the appearance of the Pitchoune neo-condensate in -rDNA. Alternatively, the reduction in nuclear size during these developmental stages might cause a crowding effect, bringing PCH closer to the nucleolus. Changes in the volumes or molecular compositions of PCH and nucleolar condensates could also alter relative affinities or biophysical properties such as surface tension and viscosity. Future studies will reveal whether one or more of these mechanisms mediate the dynamic transitions in PCH-nucleolar association during early development.

An unexpected finding in this study was the reorganization of compacted heterochromatin into a toroid-like structure, with Pitchoune filling the core when rDNA was removed. This observation led us to propose the amphiphilic model of PCH-nucleolar organization. Pitchoune is a GC-localizing RNA helicase^39,43^, has nucleolar localization signal motifs in its N-terminal tail ^49^, and its yeast ortholog is a ribosome assembly factor ^50^, altogether highlighting its dominant nucleolar affinity. We define Pitchoune’s function as an amphiphile by demonstrating two key points: (i) it forms neo-condensates in the absence of rDNA, indicating self-association, and (ii) while Pitchoune does not stably mix with HP1a, it consistently shares an interface with HP1a in both its normal and neo-condensate forms suggesting a weak affinity for HP1a. This proximity is lost when the HP1a-interacting C-terminal PxVxL motif in Pitchoune is mutated. This gradient in interaction strengths creates a hierarchical organization and stabilizes PCH-nucleolar associations. Such a layering mechanism is distinct from the formation of nested subcompartments within the nucleolus, which are immiscible but associate due to sequential rRNA synthesis, processing and ribosome assembly ^8,9,11^, however rRNA synthesis doesn’t seem to be required for Pit-HP1a associations. While synthetic amphiphiles have been shown to generate multiphasic condensates both *in vitro* ^38^ and *in vivo* within stress granules ^51^, we demonstrate this mechanism in a natural context with PCH and nucleoli and propose a generalizable role for amphiphilic molecules in co-organizing immiscible condensates in cells.

How does Pitchoune compare to other molecules required for PCH clustering around the nucleolus? For instance, depleting the Nucleoplasmin homolog NLP, CTCF, or Modulo in *Drosophila* cultured cells ^52^ or NPM1 in human and mouse cell lines ^53^ causes the loss of heterochromatin clustering around the nucleolar periphery. However, it is unclear if these proteins play a direct role in PCH-nucleolar interactions or are required to form an intact granular component that recruits other key factors. Similarly, loss of the surrounded configuration upon depletion of Pitchoune protein in cultured cells or larval eye discs could result from disrupting the composition or function of the granular component. However, mutating only the key residues in Pitchoune’s HP1a binding motif is sufficient to dissociate PCH from the nucleolus, demonstrating that Pitchoune is directly responsible for binding to HP1a and necessary for PCH-nucleolar associations. Nevertheless, a network of structural and regulatory molecules may contribute to Pitchoune localization and the overall affinity of PCH to the nucleolar edge. Proximity ligation or similar methods for identifying a complete set of molecules enriched at the PCH-nucleolar interface will help generate a full understanding of how Pitchoune promotes PCH associations and what regulates the dramatic architectural changes in early development.

The composition of condensates is determined by the multivalency and binding affinities of constituent molecules, and hierarchies in these properties can generate the higher-order structure of multi-layered condensates ^54,55^. Our findings demonstrate how disrupting condensate nucleation sites or interaction hierarchies can form new, abnormal nuclear structures or ’neocondensates’ through inherent self-associations or secondary interactions with other molecules. For example, loss of the nucleolus due to rDNA deletion leads to PCH compaction, which resembles a collapsed polymer, likely due to increased heterochromatin self-interactions. The specific responses of different components to disruption likely reflect the types and relative strengths of their encoded interaction modules. For instance, when rDNA is removed, Modulo is broadly dispersed throughout the nuclear volume, likely due to its lack of self-association and affinity for other nuclear structures. In contrast, Fibrillarin forms a separate spherical neocondensate when deprived of its processing substrate rRNA, driven by strong self-association, and Pitchoune, with strong self-association and a weak HP1a interaction motif, forms a spherical neocondensate within the compacted PCH. Given the large number of multi-component condensates within cells, it is crucial to assess whether new condensates arise when perturbing interactions (e.g., by mutating a protein binding domain). Such neomorphic responses, rather than the mutated protein itself, may cause defective cellular phenotypes or behaviors. Understanding these outcomes will be important during stress responses, aging, and cellular senescence when new condensates often form or the composition of existing condensates changes ^56^.

## Methods

### *Drosophila* Stocks and Genetics

All crosses were maintained at 25° C. To visualize the dynamics of HP1a and nucleolar assembly, live embryos from *RFP-HP1a; GFP-Fib*, *RFP-Fib; GFP-HP1a* and *RFP-HP1a; GFP-Mod* stocks were imaged. Embryos lacking rDNA were obtained as described in Falahati et al. 2016 by crossing C(1)DX/Y; *RFP-HP1a; GFP-Fib* or C(1)DX/Y; *RFP-HP1a; Pit-GFP* or C(1)DX/Y; *RFP-Fib; Pit-GFP* virgins to C(1;Y)6,y[1]w[1]f[1]/0 males. 1/4^th^ of the resulting embryos from this cross lack rDNA, and -rDNA embryos were selected based on the presence of Fibrillarin neocondensates in live and fixed embryos and DAPI morphology in fixed embryos. To knockdown Pitchoune in eye discs, eyeless-GAL4 virgin females were crossed with Pitchoune RNAi VAL20 males, and eye discs were dissected from F1 third instar larvae.

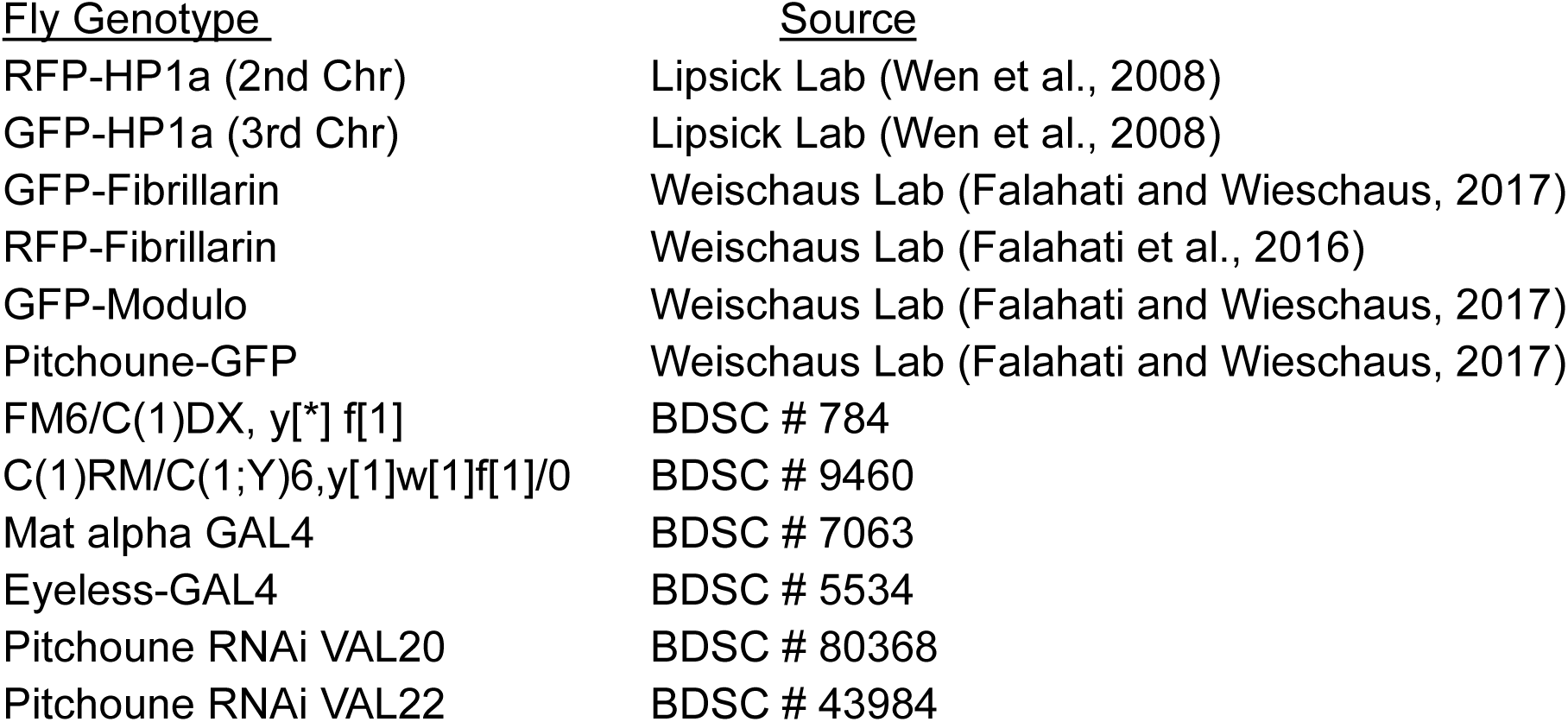

| <u>Fly Genotype</u> | <u>Source</u> |
| --- | --- |
| RFP-HP1a (2nd Chr) | Lipsick Lab (Wen et al., 2008) |
| GFP-HP1a (3rd Chr) | Lipsick Lab (Wen et al., 2008) |
| GFP-Fibrillarin | Weischaus Lab (Falahati and Wieschaus, 2017) |
| RFP-Fibrillarin | Weischaus Lab (Falahati et al., 2016) |
| GFP-Modulo | Weischaus Lab (Falahati and Wieschaus, 2017) |
| Pitchoune-GFP | Weischaus Lab (Falahati and Wieschaus, 2017) |
| FM6/C(1)DX, y[*] f[1] | BDSC # 784 |
| C(1)RM/C(1;Y)6,y[1]w[1]f[1]/0 | BDSC # 9460 |
| Mat alpha GAL4 | BDSC # 7063 |
| Eyeless-GAL4 | BDSC # 5534 |
| Pitchoune RNAi VAL20 | BDSC # 80368 |
| Pitchoune RNAi VAL22 | BDSC # 43984 |

### Preparing *Drosophila* embryos for live imaging

To collect *Drosophila* embryos for live imaging, males and females of the desired genotype were added to a plastic cage covered with apple juice agar plates and left for at least 3 days at 25°C. On the day of imaging, a fresh plate was added to the cage, and embryos were collected for two hours. After removing the plate, embryos were coated with Halocarbon oil 27 (Sigma) for staging using a dissection scope with transillumination. Stage-selected embryos were placed on a 2 x 2 inch paper towel square and dechorionated in 50% bleach for 1 min. Bleach was wicked off with a Kimwipe after 1 min, the square was washed with a small amount of distilled water, and excess water was wicked off. Dechorionated embryos were secured onto a semipermeable membrane (Lumox film, Starstedt) on a membrane slide holder using Heptane Glue. These embryos were mounted in Halocarbon oil 27 (Sigma) between the membrane and a coverslip.

### Immunostaining

Embryos were collected on apple juice-agar plates and aged till the appropriate stage, dechorionated in 50% bleach, fixed in 1:1 heptane:4% formaldehyde (Sigma) in 1XPBS for 25 mins, devitellinized in a 1:1 mixture of methanol:heptane, and stored at -20°C in methanol. Embryos were rehydrated by washing in 1xPBS+0.2% Triton (PBT). Dissected eye-discs were fixed in 4% formaldehyde in PBS for 20min, and the fixative was washed with PBT. Following washes with PBT, tissues were blocked for ∼1hr with 2% BSA in PBT, then incubated with the primary antibody at 4°C overnight. After incubation with the appropriate secondary antibody at room temperature for 2 hrs, samples were stained in DAPI and mounted onto a slide using VectaShield (Vector Laboratories) mounting medium. All primary antibodies were used at 1:250 dilution and secondaries at 1:1000. Primary antibodies: Rabbit anti-Fibrillarin (Abcam ab5821), Mouse anti-H3K9me2 (Abcam ab1220), Rabbit anti-H3K9me3 (Abcam ab8898), Mouse anti-Modulo (Gift from Mellone Lab, from Chin-Chi Chen et al., 2012), Mouse anti-Lamin, Dm0 (DSHB ADL67.10). Secondary antibodies: Goat-anti-Mouse, Alexa Fluor 488 (Invitrogen A-11001), Goat-anti-Mouse, Alexa Fluor 568 (Invitrogen A-11004), Goat-anti-Rabbit, Alexa Fluor 488 (Invitrogen A-11034), Donkey-anti-Rabbit, Alexa Fluor 568 (Invitrogen A-10042).

### DNA Fluorescent in situ hybridization (FISH) and combined Immuno-FISH

#### Probe Labelling

A probe for ITS-1 rDNA was prepared by amplifying an ∼800 bp fragment by PCR from genomic DNA using the primers 5’-ACGGTTGTTTCGCAAAAGTT-3’ and 5’-TGTTGCGAAATGTCTTAGTTTCA-3’, cloned into a pGEM T-Easy vector (Promega) and used as a template for probe synthesis. ITS-1 rDNA probe was labeled with Alexa 488, Alexa 555, or Alexa 648 using the FISH Tag DNA Multicolor Kit (ThermoFisher) following the manufacturer’s protocol. Locked nucleic acid (LNA) oligonucleotides (Integrated DNA technologies) conjugated with Cy5 or FAM were used as probes for 359bp, 1.686, and AAGAG satellite DNA repeats.

#### Hybridization

Embryos were collected on apple juice-agar plates and aged till the appropriate stage, dechorionated in 50% bleach, then fixed in 1:1 heptane: 4% formaldehyde in 1XPBS for 25 mins, devitellinized in a 1:1 mixture of methanol: heptane, and stored in -20°C. Embryos were washed in 2xSSC-T (2xSSC containing 0.1% Tween-20) with increasing formamide concentrations (20%, 40%, then 50%) for 15 min each. 100ng of DNA probes in 40 μl of hybridization solution (50% formamide, 3× SSCT, 10% dextran sulfate) was added, denatured together with the embryos at 95°C for 5 min and incubated overnight at 37°C. Following hybridization, embryos were washed twice in 2xSSCT for 30 mins at 37°C and thrice in PBT for 5 mins at room temperature. After completing washes, embryos were stained in DAPI and mounted onto a slide using VectaShield (Vector Laboratories) mounting medium.

#### Combined Immuno-FISH

Immunofluorescence was performed first on embryos for combined in situ detection of proteins and DNA sequences. Embryos were post-fixed in 4% formaldehyde for 25 mins, then processed for FISH.

### Pan-Protein Staining using 488 NHS Ester

Formaldehyde-fixed embryos were devitellinized in a 1:1 mixture of methanol: heptane and stored in methanol. Embryos were rehydrated by washing in 1xPBS+0.2% Triton (PBT). After washing off methanol, embryos were stained in the diluted Atto 488 NHS ester fluorophore (Sigma) (1:50) from a 10mg/ml stock in 0.1% PBST for 6 h at 4 °C followed by washing in PBT three times for 30 mins each at room temperature. Embryos were stained in DAPI for ten minutes and mounted onto a slide using VectaShield (Vector Laboratories) mounting medium.

### Propidium Iodide Staining

Formaldehyde-fixed and Heptane devitellinized embryos were rehydrated by washing in 1xPBS+0.2% Triton (PBT). Samples were equilibrated in 2X SSC. RNase-treated controls alone were incubated in 100 μg/mL DNase-free RNase in 2X SSC for 20 minutes at 37°C. After washing away the RNase with 2X SSC, embryos were incubated in 500nM of Propidium Iodide (Invitrogen) in 2X SSC for 10 mins at room temperature. Samples were rinsed in 2X SSC, stained with DAPI, and mounted on a slide with VectaShield (Vector Laboratories) mounting medium.

### Microscopy

Imaging was performed on a Zeiss LSM880 Airy Scan microscope (Airy Fast mode) with a 63X NA 1.4 oil immersion objective at room temperature. Depending on the fluorophore, 405, 488, 514, or 633nm laser lines were used for excitation with appropriate filter sets. Laser intensity values, detector gain, image size, zoom, z-stack intervals, and time intervals (for time-lapse acquisitions) were adjusted to minimize bleaching and ensure uniform detection across all AiryScan detection elements. Once standardized for an experiment, settings were kept identical across all samples in the experimental groups. Raw images were processed using Zeiss ZEN Black software with the AiryScan processing module for reconstruction and subsequent image analysis.

### Modeling

To better understand the association of PCH with the nucleolus, we developed a physical model that simulates the interactions between different types of molecules found in these biomolecular condensates. In our physical model, we simulate four components of the nucleus: PCH (H) and ribosomal DNA (rD) as long polymers, and Fibrillarin (F) and an amphiphilic protein (X) as independent, single monomers, which we hereafter refer to as beads. Since the experiments focused on the PCH domain of *Drosophila*, our physical model only simulates this specific region of the genome (30% of the genome), not the entire genome. This allows us to study the dynamics of this particular region of the genome more accurately and efficiently. PCH is modeled as a semiflexible bead-spring polymer chain in which N beads are connected by N-1 harmonic springs. Each bead of the chain represents a cluster of PCH containing approximately 5 kilo base pairs of DNA, with a diameter of approximately 30 nm. The semi-flexibility of the chain is determined by its persistence length, which is taken to be 60 nm (2 beads) in accordance with previous studies that indicate chromatin has a persistence length between 50 to 100 nm (Wachsmuth et al. 2016). We represent rDNA as a self-avoiding chain that occupies approximately 20% of the middle domain of PCH. Fibrillarin and the amphiphilic protein X are modeled using single, diffusive beads where each protein has distinct interactions with the polymer and other proteins.

To simplify the model, we assume that the size of each protein bead is equal to the size of the heterochromatin bead, with both having a diameter *σ*. The non-bonding interactions between polymer-polymer, protein-protein, and polymer-protein are modeled using a standard Lennard-Jones (LJ) potential. The LJ potential is truncated at the distance 2.5 *σ*, meaning that the interaction between the beads is only non-zero if they are within 2.5 *σ* distance. At very short distances between the beads, the LJ potential is strongly repulsive (representing the excluded volume of two molecules). At intermediate spacings, the LJ potential is attractive, with a strength adjusted to model the different states of chromatin, such as its compaction or decondensation and the phase-separating tendency of the proteins. In our model, the polymer and the proteins are confined within a spherical boundary that represents the nucleus of the cell. This boundary mimics the effect of the nuclear envelope, which constrains the movement of these beads and affects their interactions with each other. In our study, we used the LAMMPS (Large-scale Atomic/Molecular Massively Parallel Simulator) package to simulate the behavior of our biomolecular system (Thompson et al. 2022). LAMMPS uses Brownian dynamics, which accounts for the viscous forces acting on the beads, and a stochastic force (Langevin thermostat) to ensure that the system of beads and solvent is maintained at a constant temperature (NVT ensemble). This allows us to model the interactions between polymer-polymer, polymer-protein, and protein-protein beads accurately and study the behavior of the system over time.

### Rationale for the choice of parameters in the coarse-grained model

To analyze the experimental observations of phase separation in the nucleus, we study a minimal model with four crucial components: PCH (H), rDNA (rD), Fibrillarin (F), and an amphiphilic protein (X) that binds both nucleolar and PCH components. The parameters in the simulations include the number of molecules of each component α (*N*_*α*_), the strength of the attractions between two components (denoted by indices *β* and *γ*, *∊*_*βγ*;_), and the size of the confinement (*R*_*c*_). The fraction of the nucleus that is hydrated (does not contain PCH or the other proteins) is obtained from the relative difference between the confinement volume and the volumes of PCH and the other proteins.

#### Bonding potential between monomers in the polymer made of PCH and rDNA

Adjacent beads on the polymer chain are interconnected by harmonic springs using the potential function:

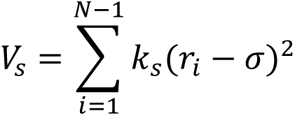

Here, *r*_*i*_ represents the distance between the *i*-th and (*i* + 1)-th beads. The spring constant and equilibrium distance between neighboring beads are denoted as *k*_*s*_ and *σ* respectively. In our simulations, the spring constant *k*_*s*_ is set to 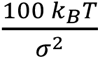 to ensure the presence of rigid bonds between adjacent beads of the polymer chain.

#### Attraction strength (*∊*_*βγ*_)

The Lennard-Jones potential is used to model the attraction between any two non-bonded beads:

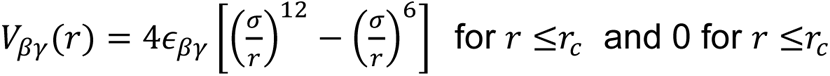

Here, the symbol *∊*_*βγ*_ represents the attraction strength between beads of type *β* and *γ*, where *β*, *γ* ∈ {*H*, *r*D, *F*, *X*} represent PCH, rDNA, Fibrillarin, and amphiphilic protein, respectively. For instance, *∊*_*FX*_ represents the attraction strength between Fibrillarin and amphiphilic protein beads. When dealing with attractive interactions between chromatin-chromatin and chromatin-protein beads, a distance cutoff of *r*_*c*_ = 2.5*σ* is used for the Lennard-Jones potential, beyond which the interaction is set to zero. To account for only excluded volume interactions (with no attractions) using the same potential, a cutoff of *r*_*c*_ = 2^1/6^*σ* and *∊* = 1*k*_*B*_*T* are employed. This choice is made because the potential energy is at its minimum at that point, and the resulting force on a bead is zero. As there are four components in our model, there are a total of 10 combinations (n(n+1)/2), where n is number of components, of attraction strength parameters between the different components.

#### Confinement size

The size of the confinement is determined by defining the volume fraction of chromatin *ϕ*:

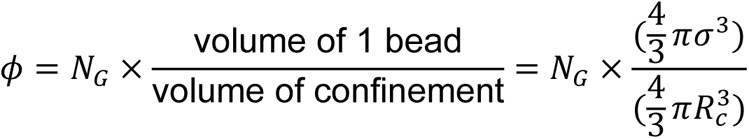

Here, *N*_*G*_ represents the total number of beads in the *Drosophila* genome, where each bead corresponds to 5 kilobase pairs (kbps) of DNA. The diameter of a spherical bead, denoted as *σ*, is taken to be 30 nm (see ^59^ for further explanation). The total length of the diploid *Drosophila* genome is 360 Mbps, and *N_G_* can be calculated by dividing the *Drosophila* genome length by the amount of DNA represented by one bead (5 kbps). The volume fraction of chromatin (*ϕ*) is commonly reported as ∼ 0.1 in existing literature (Qi & Zhang, 2021; Tripathi & Menon, 2019). Using the equation above, the calculated radius of the confinement (*R*_*c*_) is found to be 45*σ*.

#### Number of molecules in PCH and rDNA

We model the PCH domain as a polymer chain composed of *N* = 10,000 beads, which represents approximately 30% of the beads in the entire genome. Among these 10,000 beads, 20% are designated as rDNA, resulting in a total of 4,000 rDNA beads. We do not explicitly model the rest of the genome since the experiments show that the nucleolar components are localized near PCH.

#### Concentration of Fibrillarin and amphiphilic protein

The concentration of Fibrillarin and amphiphilic protein is calculated as follows:

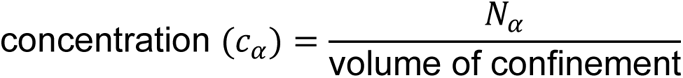

Here, the symbol *α* represents the type of protein, where *c*_*α*_= *c*_*F*_ for the fibrillarin concentration and *c*_*α*_ = *c*_*X*_ for the amphiphilic protein concentration. After defining the parameters and obtaining the value for the confinement radius parameter from the assumed volume fraction of PCH, we proceed with an initial simulation, focusing on a single component, namely the Fibrillarin protein, within the confinement. During this simulation, we vary the concentration (*c*_*F*_) and the attraction strength (*∊*_*FF*_) of Fibrillarin. Our experimental results demonstrate that Fibrillarin undergoes phase separation independently of the other components (rDNA or PCH) during cycle 14 (refer to Figure 2A). These initial-stage simulation results yield a phase diagram, which indicates that a minimum attraction strength of 1.3 − 2.0*k*_B_T is required to condense fibrillarin particles within the concentration range of *c*_*F*_ = 0.0013 − 0.013. The reported concentration of nucleolar particles is *c* = 0.015 (Qi & Zhang, 2021). Consequently, we maintain a fixed concentration of Fibrillarin at *c*_*F*_ = 0.013 when simulating all the other protein components (**Extended Data Fig. 4a**). Since the concentration of amphiphilic protein is not determined from the experiments, we conducted multiple simulations, systematically varying the amphiphilic protein concentration (*c*_*X*_) within the range of 0.0013 − 0.13. The results of these simulations are discussed in **Fig. 6a-b**.

#### Parameter range of *∊*_*HH*_

To understand the behavior of each component separately (before including interactions between different components) in the next stage, we conducted simulations specifically focusing only on the PCH chain. The PCH chain represents a condensed chromatin region within the nucleus which implies self-attractive interactions of the beads representing the polymer. During these simulations, we varied the attraction strength between PCH beads (*∊*_*HH*_) from 0 to 0.5*kk*_B_T. Our results revealed that within the range of *∊*_*HH*_ = 0.35 − 0.5 *k*_B_T (**Extended Data Fig. 4b**), the PCH chain underwent collapse, resulting in a condensed conformation that is phase-separated from the aqueous component of the system (not simulated explicitly).

#### Parameter value of *∊*_*r*D−*F*_

We next simulate the self-organization due to the interactions between the three components (where each component so far was considered alone): PCH, rDNA, and Fibrillarin. We set *∊*_*HH*_= 0.35 *k*_B_T and *∊*_*FF*_= 2 *k*_B_T, incorporating only excluded-volume interactions (hard-core, repulsive interactions) between H-F, i.e., there is no direct attraction between PCH and Fibrillarin as implied by the experiments. By varying the attraction strength between rDNA and Fibrillarin (*∊*_*r*D−*F*_), we made the following observations based on our simulation results: a weaker attraction (and *∊*_*r*D−*F*_ = 0.75 *k*_B_T) resulted in rDNA wrapping around the condensed Fibrillarin phase, while a stronger attraction (and *∊*_*r*D−*F*_= 2 *k*_B_T) led to the condensation of rDNA within the Fibrillarin complex. This latter observation aligns with our experimental findings. Therefore, we select *∊*_*r*D−*F*_ = 2 *k*_B_T as the parameter value in the subsequent simulations (**Extended Data Fig. 4c**).

#### Parameter ranges of *∊*_*FF*_ and *∊*_*XX*_

Finally, we introduce the fourth component, an amphiphilic protein ’X’, which we suggest may interact attractively with both PCH and Fibrillarin. Initially, we investigated the relative attraction strengths between Fibrillarin and protein X when considering the same concentration for both. We explore all possible combinations of *∊*_*FF*_ both greater than and less than *∊*_*XX*_. For *∊*_*XX*_ ≥ *∊*_*FF*_, we observe PCH surrounding the amphiphilic protein-rich phase, but just a partial wetting between the amphiphilic protein-rich phase and the Fibrillarin-rich phase (**Extended Data Fig. 4d**). Only when *∊*_*XX*<_*∊*_*FF*_ do we observe PCH surrounding the amphiphilic protein-rich phase, which in turn surrounds the Fibrillarin condensate, consistent with the experimental results in wild type embryos.

#### Parameter range of *∊*_*FX*_

We proceeded to vary the attraction strength between Fibrillarin and amphiphilic protein X within the range of *∊*_*FX*_ = 1.5*k*_B_T. When the attraction strength is relatively low (*∊*_*FX*_ ≤ 0.75*k*_B_T), the fibrillarin-rich phase and the amphiphilic protein-rich phase do not associate with each other. At moderate attraction strengths (*∊*_*FX*_ ≥ 1*k*_B_T and (*∊*_*FX*_ ≤ 1.25*k*_B_T), the Fibrillarin-rich and amphiphilic protein-rich phases partially wet each other. Finally, at higher attraction strengths (*∊*_*FX*_= 1.5*k*_B_T), the amphiphilic protein completely wets Fibrillarin (**Extended Data Fig. 4e-f**).

#### Parameter range of *c*_*X*_

Additionally, we explored variations in the concentration of amphiphilic protein X (*c*_*X*_). Notably, for higher concentrations of protein X (*c*_*X*_ = 0.005 − 0.013), we observed that heterochromatin tends to completely engulf the amphiphilic protein-rich phase, which in turn engulfs Fibrillarin (**Fig. 6a**).

### Cell culture

*Drosophila* S2 cells were cultured in Schneider’s *Drosophila* Medium (Gibco) with 10% FBS and 1% antibiotic-antimycotic (Gibco) at 25°C. For transfections, cells were seeded at 5 x 10^5^ cells/mL in 6-well plates 24 hours prior. 1 µg of plasmid DNA was diluted in 100 µL of serum-free medium and mixed with 2 µL of TransIT-2020 Transfection Reagent (Mirus Bio). After a 15-min incubation, the DNA-reagent complexes were added to the cells and incubated at 25°C for 48-72 hours before visualizing.

### Plasmids/Recombinant DNA

Codon-optimized gene blocks for Pitchoune, Fibrillarin, Modulo, Polr1E, and HP1a were synthesized by Twist Biosciences and cloned into pCOPIA vectors fused with fluorescent protein tags. Site-directed mutagenesis was performed to introduce PxVxL and DQVD mutations into the full-length Pitchoune using the Q5® Site-Directed Mutagenesis Kit (New England Biolabs) using these primers:

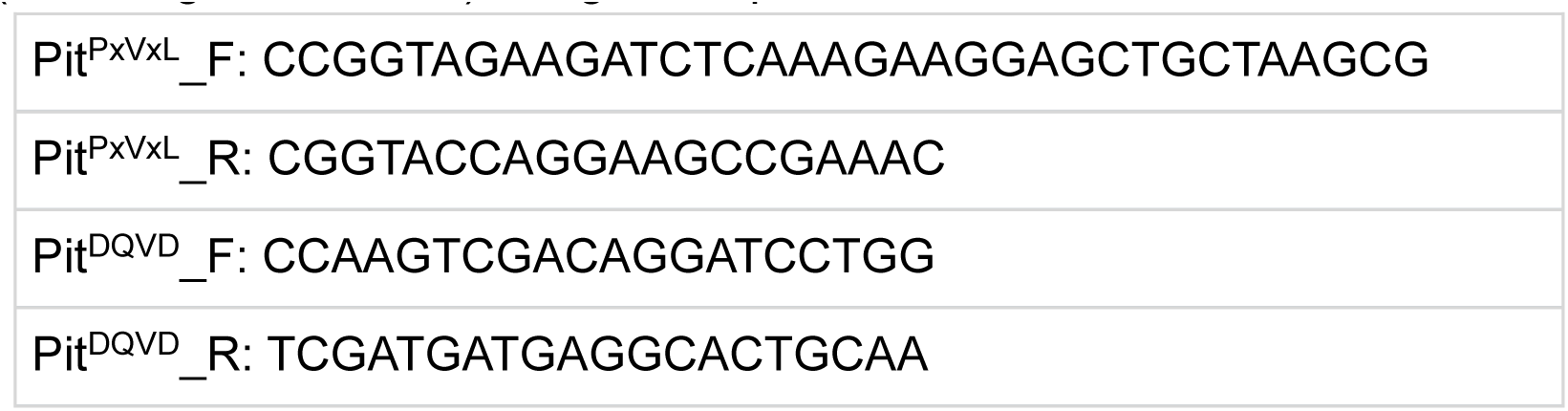

### Pitchoune RNAi

Genomic DNA from S2 cells was used as the template for PCR amplification with the primers listed below to generate the amplicon for Pitchoune RNAi targeting its 3’ untranslated region (3’UTR). Mock RNAi targeting the y gene was used as a negative control.

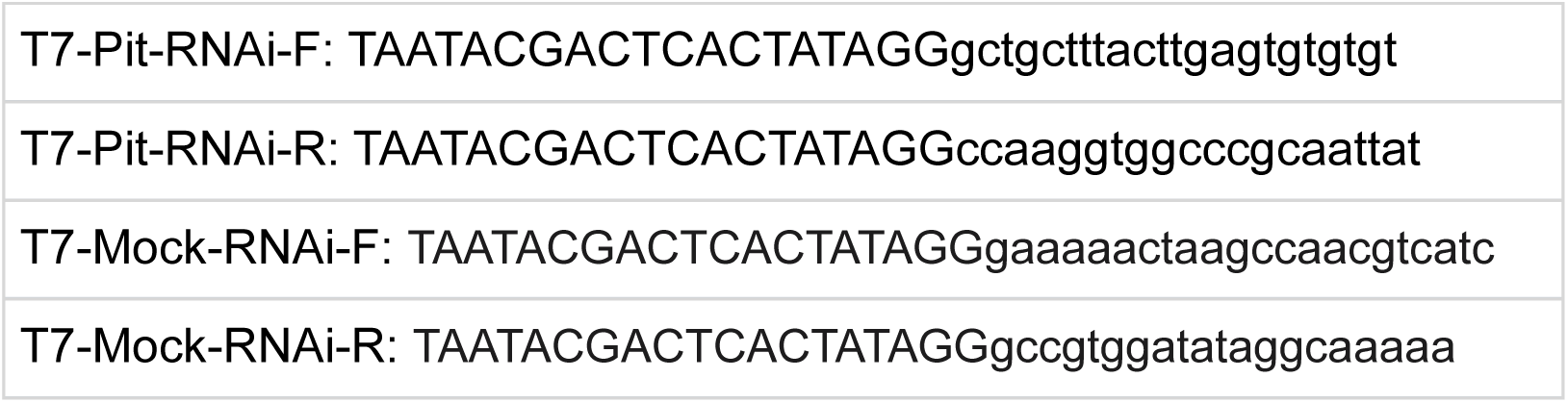

Hi-Scribe T7 Synthesis kit (New England Biolabs) was used to synthesize double-stranded RNA (dsRNA) using manufacturer’s protocol. Following synthesis, RNA purification was carried out utilizing the MinElute RNeasy Kit (Qiagen). The purified RNA was then diluted to 1 μg/μl. For the RNAi experiment, 3-5 μg of dsRNA with DOTAP liposomal transfection reagent (Roche) was used per 0.5 x 10^6 cells. Cells were analyzed 5 or 6 days after the initiation of RNAi.

### Total RNA Preparation, cDNA Synthesis, and Quantitative PCR

Total RNA from S2R+ cells was extracted by homogenizing in TRIzol Reagent (Invitrogen). 0.5 volume of chloroform was added, the mixture was shaken for 15 seconds, incubated for 3 minutes, and centrifuged at 12,000 x g for 15 minutes at 4°C. The aqueous phase was transferred to a new tube, mixed with 500 µL of isopropanol, incubated for 10 minutes, and centrifuged at 12,000 x g for 10 minutes at 4°C. The RNA pellet was washed with 75% ethanol, centrifuged at 7,500 x g for 5 minutes at 4°C, air-dried for 5-10 minutes, and resuspended in RNase-free water. Total RNA was treated with DNA-free DNA removal kit (Invitrogen) per manufacturer’s protocol to remove any contaminating genomic DNA. RNA was converted to cDNA with GoScript Reverse Transcriptase Kit (Promega) using random primers, and Real-Time PCR was performed using PerfeCTa SYBR^®^ Green FastMix (Quantabio) on the Biorad CFX96 Real-Time PCR Detection system. Analysis was performed using the 2^−ΔΔCt^ method, with relative mRNA levels of Pitchoune normalized to β-actin.

### Quantitative Image Analysis

3D measurements were performed using Arivis Vision4D (Zeiss) while ImageJ was used for all 2D measurements. The details of each analysis pipeline used in this study are listed below:

#### Measuring the fraction of the nucleolar edge occupied by HP1a or H3K9me2

<u>Fig. 1c:</u> Nuclei were manually chosen for analyses ∼15-30 mins into the specified interphase and defined as "Early Cycle ", while those observed from ∼50-70 mins were defined as "Late Cycle ". Preprocessing steps include background subtraction and denoising using Gaussian blur. HP1a and Fibrillarin were segmented in 3D using the "Intensity Threshold Segmenter" with the Auto segmentation method. To calculate HP1a occupancy at the nucleolar edge, the nucleolus was dilated by 1 pixel and subtracted from the dilated object to create a 1-pixel shell around the nucleolus. The nucleolus shell was intersected with HP1a segments to calculate the volume fraction of the nucleolar shell that overlaps with HP1a.

<u>Fig. 6c:</u> Same as above, except the nucleolus was dilated by 2 pixels to create a 2-pixel shell around the nucleolus. The shell was intersected with H3K9me2 segments in nuclei from eye-discs to calculate the fraction of the nucleolar shell that overlaps with H3K9me2.

<u>Fig. 6e:</u> Same as above, except the nucleolus was dilated by 4 pixels to generate a 4-pixel shell around the nucleolus. The thickness of the nucleolar shell scaled with nucleolar size in the different cell types.

#### Measuring Distances

<u>Fig. 2b:</u> HP1a and Fibrillarin were segmented using the "Intensity Threshold Segmenter" with the Auto segmentation method. The distance between HP1a and its nearest Fibrillarin segment was measured in 3D using the "Distances" feature in Arivis.

<u>Fig. 2f:</u> AAGAG and 1.686 were segmented using the "Intensity Threshold Segmenter" with the Auto segmentation method. The distance between AAGAG and its nearest 1.686 segment was measured in 3D using the "Distances" feature in Arivis.

<u>Extended Data Fig. 2e:</u> Preprocessing steps include background subtraction and denoising using Gaussian blur. 1.686 or 359bp foci and Fibrillarin were manually segmented using the "Intensity Threshold Segmenter" with the Simple segmentation method. The distance between a Fibrillarin segment and its nearest 1.686 or 359bp locus was measured in 3D using the "Distances" feature in Arivis.

#### Measuring Aspect Ratio of HP1a

<u>Fig. 2d:</u> Individual nuclei were manually selected 15 mins after the start of Cycle 15. Preprocessing steps include background subtraction and denoising using Gaussian blur. HP1a was segmented using Auto thresholding using Otsu. The aspect ratio of the segment was determined using the Analyze Particles feature in Fiji.

#### Line Scans

<u>Fig. 6g:</u> Nucleoli were segmented in Fiji using Otsu’s method. The Feret’s diameter was calculated for each nucleolus, and intensity values were measured along the Feret’s diameter. Intensities for each profile were normalized to its average value. The Feret’s diameter was normalized by setting its range from 0 to 1.

#### Pitchoune neo-condensate formation measurements

<u>Fig. 5d-f:</u> To measure the dynamics of the formation of Pitchoune in -rDNA embryos, maximum intensity projections of Amnioserosa were first preprocessed in Fiji using Subtract background and a Gaussian Blur filter. Auto thresholding for each time point was performed using the Yen method. Using Analyze Particles, the area, circularity, and mean intensity of each segment of Pitchoune was extracted. The mean intensity over time was normalized to its value at T=0.

### Statistical Analysis

Data were plotted, and statistical analyses were performed using GraphPad Prism8. P-values were calculated using unpaired two-tailed t-tests.

## Data availability statement

All data supporting the results of this study are included in the manuscript. Reagents used in this study are available upon request.

## Code availability statement

For the simulation component of this study, all simulations, analyses, and visualizations were conducted using publicly available software packages and custom-developed codes. Langevin Dynamics simulations were performed using LAMMPS (version 23 June 2022), and visualizations were generated using Ovito (version 3.7.11). The complete set of codes required to reproduce the simulations is available in the following repository: https://github.com/gauravbajpaimaths/Coarse-grained_model_of_nucleolar_heterochromatin_condensates

## ACKNOWLEDGEMENTS

We thank Eric Weischaus (Princeton University) for generously sharing the fluorescently tagged nucleolar fly lines. The Karpen lab acknowledges critical support from the National Institutes of Health (R35GM139653), and SS and GHK are grateful for the support of the Volkswagen Stiftung (98196).

## AUTHOR CONTRIBUTIONS

SR: conceptualization, experimentation, data acquisition, formal analysis, validation, investigation, methodology, and writing

OA: conceptualization, investigation, formal analysis, and writing

GB: simulations, data acquisition, formal analysis, methodology, and writing

KL: experimentation and data acquisition

SC: experimentation, data acquisition, reagent preparation, and formal analysis

SS: writing, funding acquisition, and supervision

GHK: conceptualization, writing, funding acquisition, and supervision

## LEGENDS FOR SUPPLEMENTARY MOVIES

**Supplementary Movie 1:** 3D rendering of RFP-Fib (green) and GFP-HP1a (magenta) in epidermal cells of a Stage 16 Drosophila embryo.

**Supplementary Movie 2:** Live imaging of RFP-Fib (green) and GFP-HP1a (magenta) in nuclear cycle 13 of Drosophila embryogenesis.

**Supplementary Movie 3:** Live imaging of RFP-Fib (green) and GFP-HP1a (magenta) in nuclear cycle 14 of Drosophila embryogenesis.

**Supplementary Movie 4:** Live imaging of RFP-Fib (green) and GFP-HP1a (magenta) in nuclear cycle 15 of Drosophila embryogenesis.

**Supplementary Movie 5:** Live imaging of RFP-Fib (green) and GFP-HP1a (magenta) in nuclear cycle 16 of Drosophila embryogenesis.

**Supplementary Movie 6:** Live imaging of RFP-Fib (green) and GFP-HP1a (magenta) in nuclear cycle 17 of Drosophila embryogenesis.

**Supplementary Movie 7:** Live imaging of GFP-Fib (green) and RFP-HP1a (magenta) in nuclear amnioserosa nuclei in embryos lacking rDNA.

**Supplementary Movie 8:** Simulations of coarse-grained modeling of rDNA (red), Fibrillarin (green), PCH (magenta), and an amphiphilic protein (yellow) with the parameters listed in the 4X4 matrix in Fig. 4c.

**Supplementary Movie 9:** Live imaging of Pitchoune-GFP (green) and RFP-HP1a (magenta) in nuclear amnioserosa nuclei in embryos lacking rDNA.

---

**Extended Data Fig. 1:**
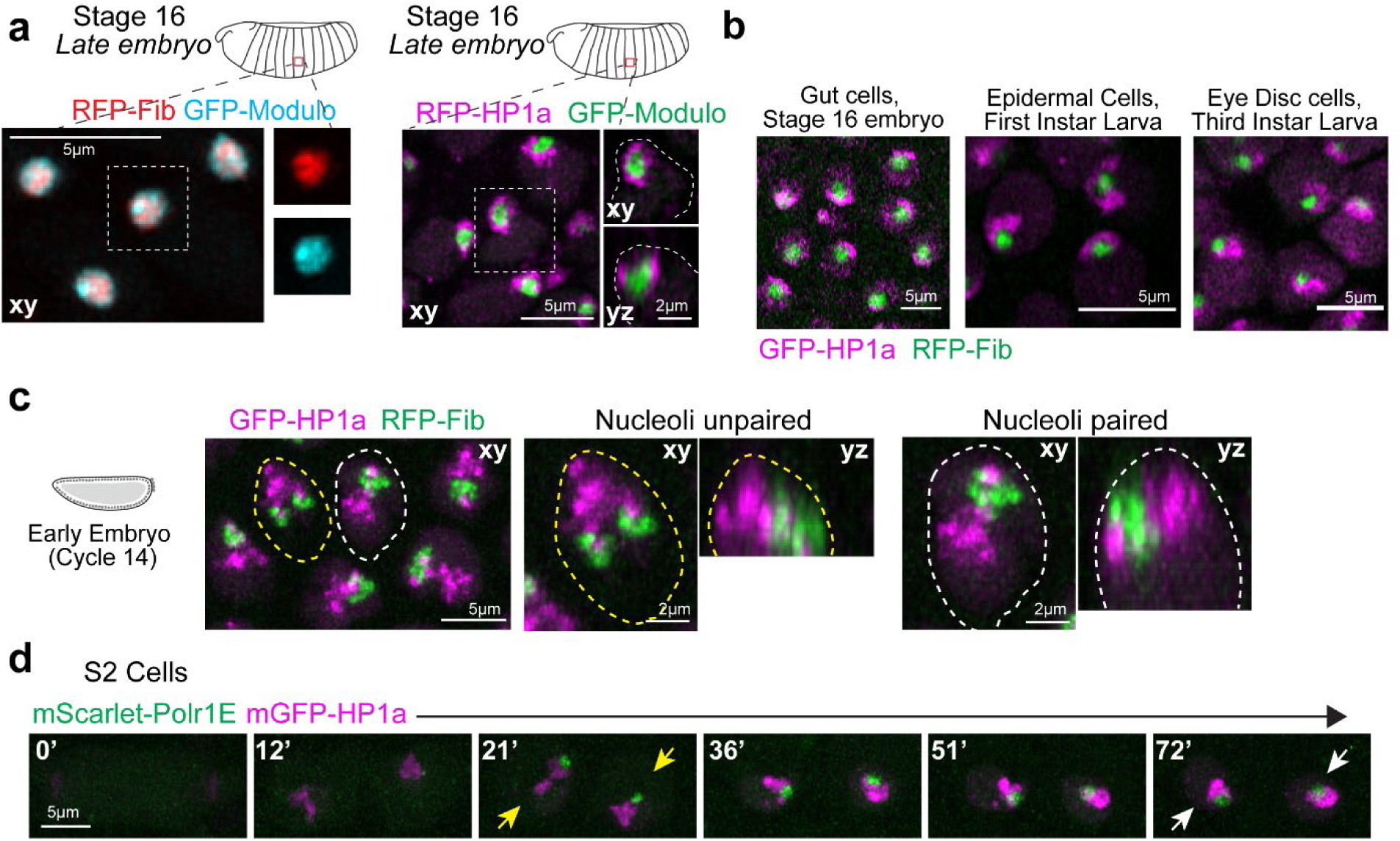
PCH organization relative to the nucleolus in Drosophila embryos, larval tissues, and cultures cell lines. (**a**) (Left) Distribution of GFP-Modulo (cyan) and RFP-Fibrillarin (red) in nucleoli of a live late-stage (Stage 16) embryo. (Right) Maximum intensity projections showing PCH localization labeled with RFP-HP1a (magenta) and nucleoli marked by GFP-Modulo (green) in live epidermal nuclei from a late-stage *Drosophila* embryo (Stage 16, ∼14-16hr). The nucleus outlined by the white dashed box is magnified and presented in xy and xz views, with white dashed lines indicating the nuclear boundary. (**b**) Stills of live nuclei expressing GFP-HP1a (magenta) and RFP-Fib (green) in gut cells from a Stage 16 embryo, first instar larval epidermal cells, and third instar larval eye disc. (**c**) Nuclei from Cycle 14 embryos expressing GFP-HP1a (magenta) and RFP-Fib (green), showing two unpaired nucleoli (yellow dashed line) and one paired nucleolus (white dashed line). Both nuclei have been enlarged to show the “extended” conformation of HP1a relative to the nucleolus in the xy and yz views. (**d**) Time-lapse stills of two daughter S2 cells exiting mitosis, transfected with mScarlet-Polr1E, a Pol-I subunit (green) and mGFP-HP1a (magenta). Numbers on the top left corner indicate time in minutes from the end of mitosis. Yellow arrows indicate HP1a in an extended conformation from the nucleolus, while white arrows indicate HP1a in the surrounded conformation.

**Extended Data Fig. 2:**
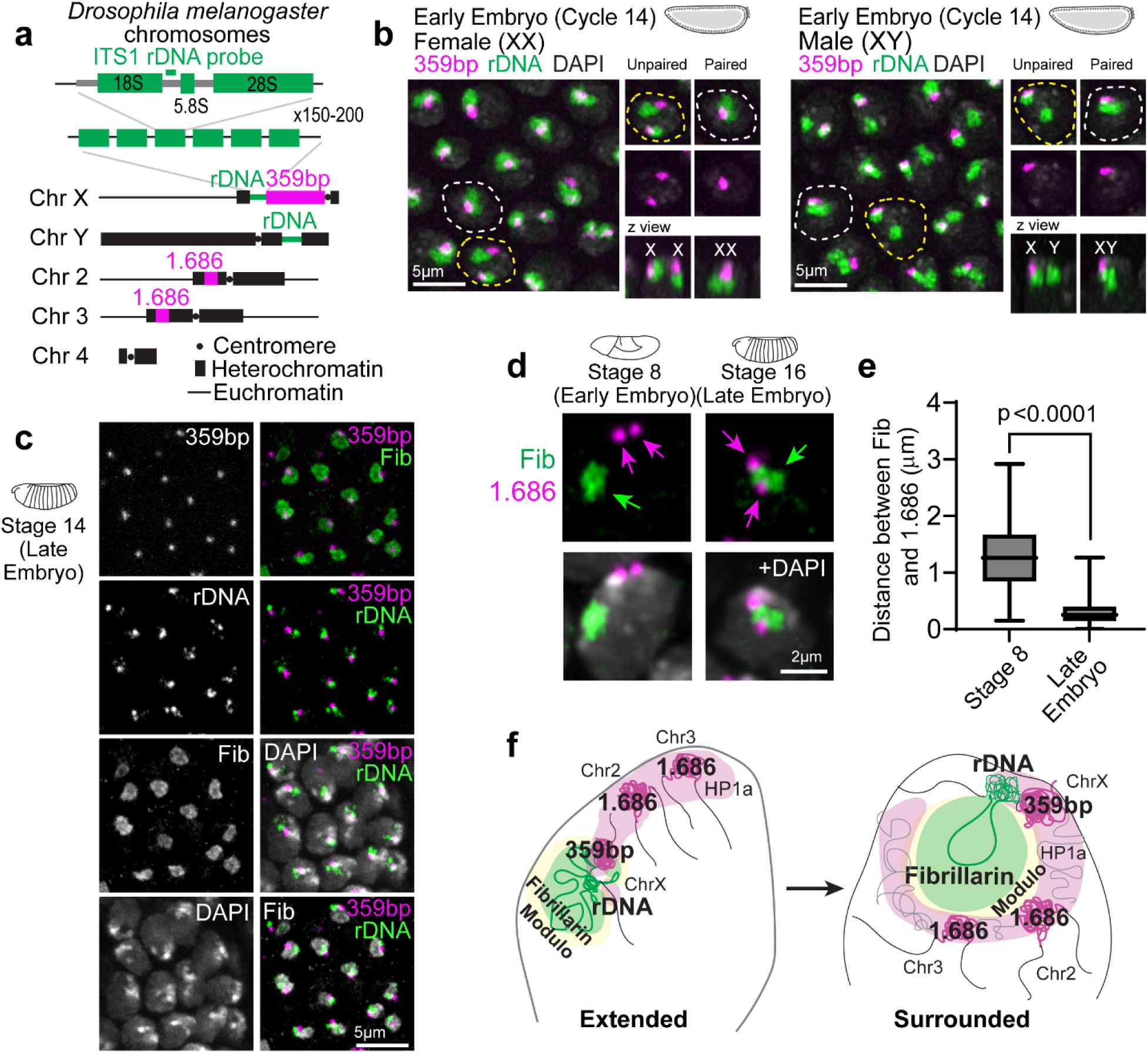
Dynamics of PCH reorganization relative to the nucleolus during *Drosophila* embryonic development. (**a**) Schematic representation of pericentromeric satellite repeats (359bp and 1.686) and rDNA repeats in *Drosophila* melanogaster chromosomes. The schematic of ribosomal DNA (rDNA) arrays indicates the position of the ITS-1 rDNA probe used for FISH. (**b**) Localization of 359bp satellite DNA (magenta) and rDNA (green) in female and male early (Cycle 14) embryos. The dashed white line indicates a nucleus with paired nucleoli, while the dashed yellow line marks a nucleus with unpaired nucleoli. The nuclear boundary is determined by DAPI staining. (**c**) Combined immuno-FISH stained for 359bp, ITS-1 rDNA, Fibrillarin, and DAPI in late embryos. (**d**) Combined immuno-FISH of 1.686 (magenta arrows), Fibrillarin (green arrow) and DAPI (grey) in a nucleus from epidermal cells of early (Stage 8, ∼Cycle 15) and late (Stage 16) *Drosophila* embryos. (**e**) Distance between the centers of geometry of 1.686 and Fibrillarin in Stage 8 (∼Cycle 15) and Stage 16 (Late) in *Drosophila* nuclei. n>60 loci (from 3 embryos) at each developmental stage. Bar graphs extend from 25^th^ to 75^th^ percentile, error bars: min to max. (**f**) Schematic summarizing the dynamic reorganization of nucleoli and PCH during *Drosophila* development, highlighting key proteins and DNA elements involved.

**Extended Data Fig. 3:**
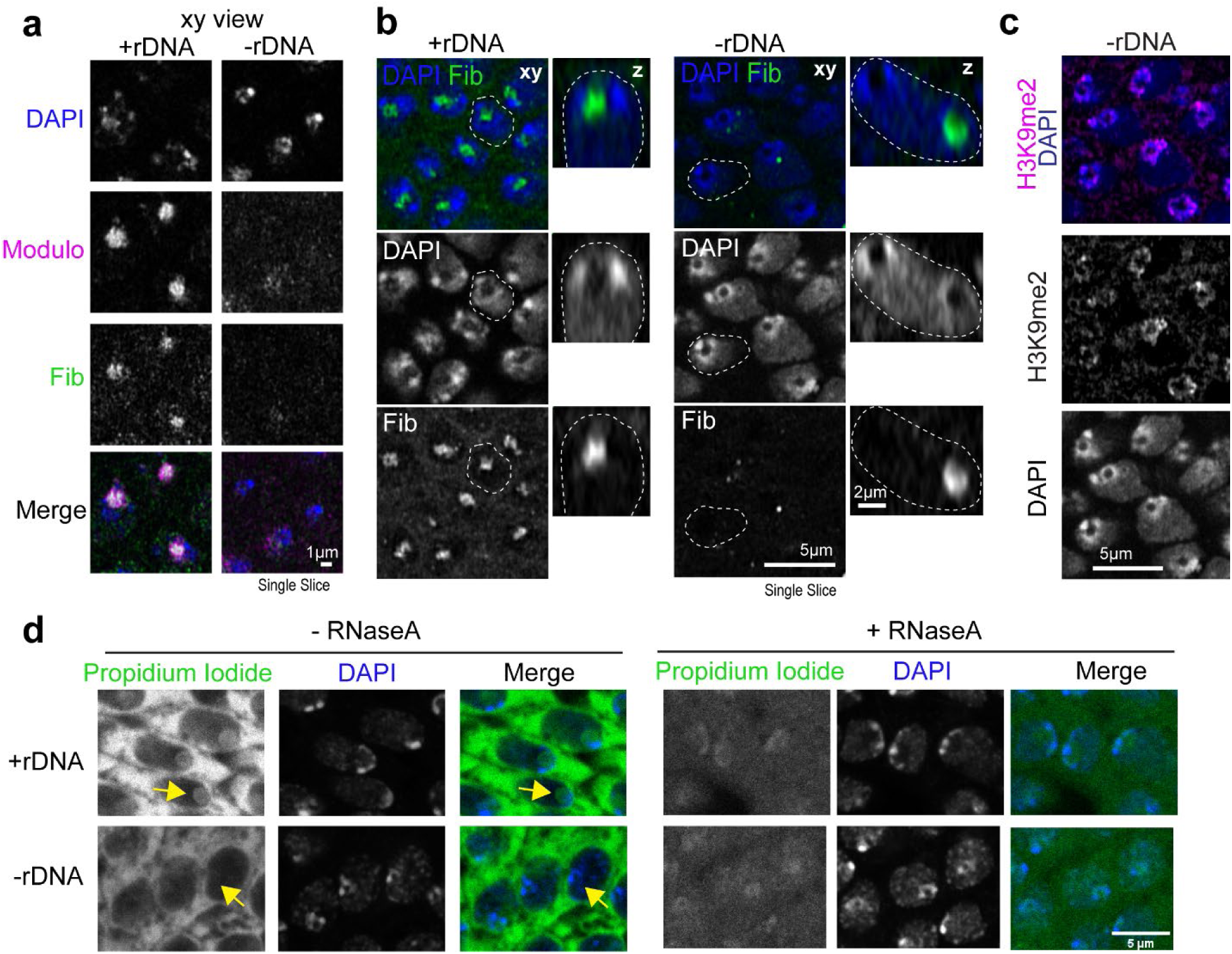
The PCH void in -rDNA embryos does not stain for DAPI, Fibrillarin, Modulo, H3K9me2 or Propidium Iodide (RNA). (**a**) Representative images of fixed nuclei from wildtype late embryos and mutant embryos lacking rDNA stained for Modulo (magenta), Fibrillarin (green) and DAPI (blue). (**b**) Left: Representative images of fixed nuclei from Stage 14-16 (late) wildtype embryos and mutant embryos lacking rDNA showing Fibrillarin (green) and DAPI (blue). Right: Nuclei marked with white dashed outlines on the left are shown in the z plane. (**c**) H3K9me2 immunofluorescence (magenta) and DAPI (blue) staining in -rDNA nuclei in Stage 14-16 (late) *Drosophila* embryos. (**d**) Representative images of fixed nuclei from Stage 14-16 (late) wildtype and mutant embryos lacking rDNA stained with Propidium Iodide (green) and DAPI (blue) without (left) and with (right) RNaseA. The yellow arrow points to the RNA staining in the nucleolus in +rDNA and lack of propodium iodide staining in the PCH void of -rDNA nuclei.

**Extended Data Fig. 4:**
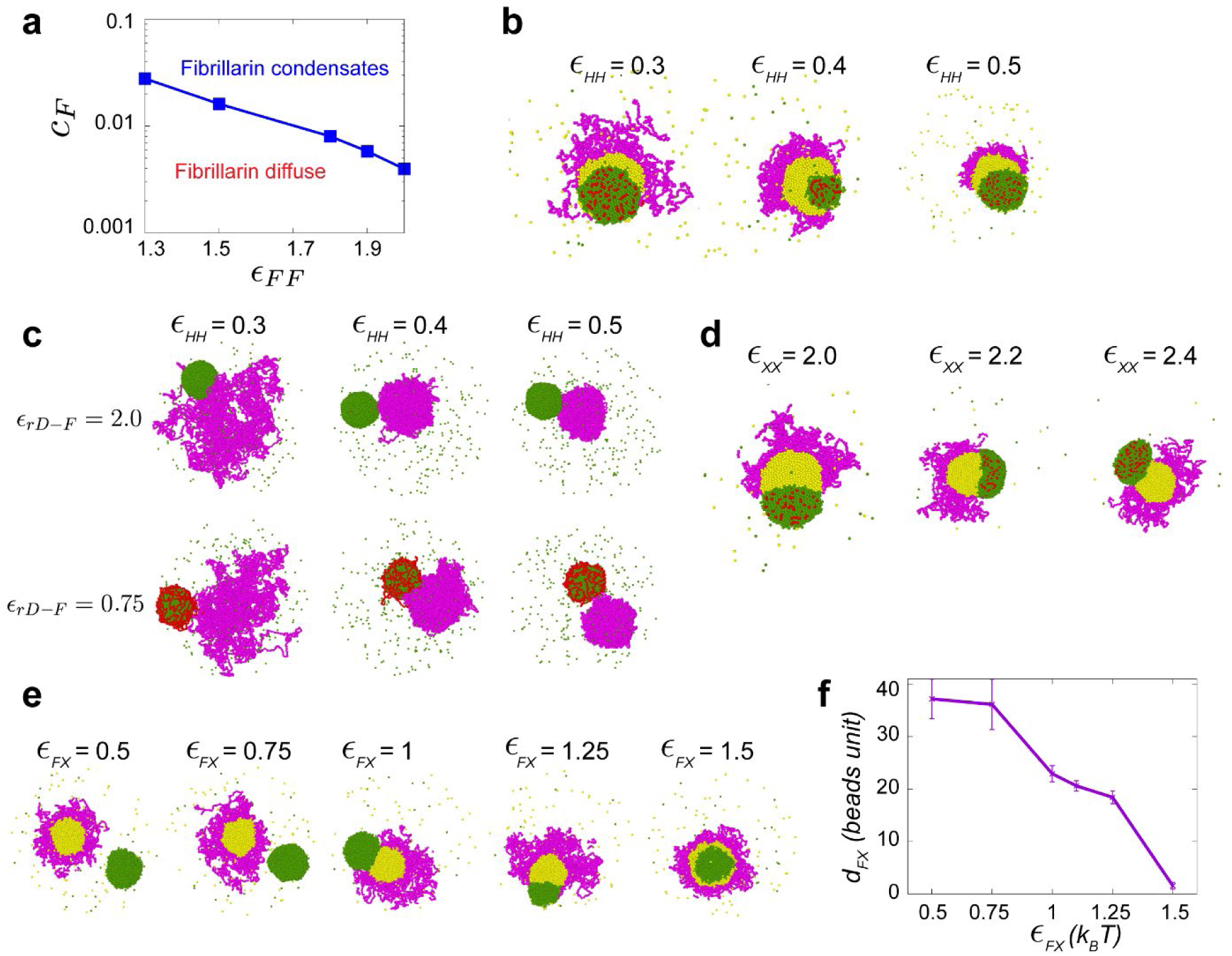
Coarse-grained model for the assembly of the nucleolus and PCH. (**a**) The phase diagram illustrates the minimum attraction strength required to condense Fibrillarin at different concentrations. (**b**) Simulation endpoint snapshots depict the outcomes of varying the attraction strengths between beads of the PCH (H) polymer chain (*ɛ_HH_*). (**c**) Simulation snapshots depict varying attraction strengths between rDNA-Fibrillarin (top to bottom) and beads of the PCH polymer chain (left to right). (**d**) Simulation endpoint snapshots depict the outcomes of varying X-X attraction strengths (*ɛ_XX_*). (**e**) Simulation endpoint snapshots depict the outcomes of varying attraction strengths between Fibrillarin and protein X (*ɛ_FX_*) in the -rDNA condition. (**f**) The average distance (*d_FX_*) between Fibrillarin and protein X condensates from their center of mass is measured for different attraction strengths. Error Bars represent s.d.

**Extended Data Fig. 5:**
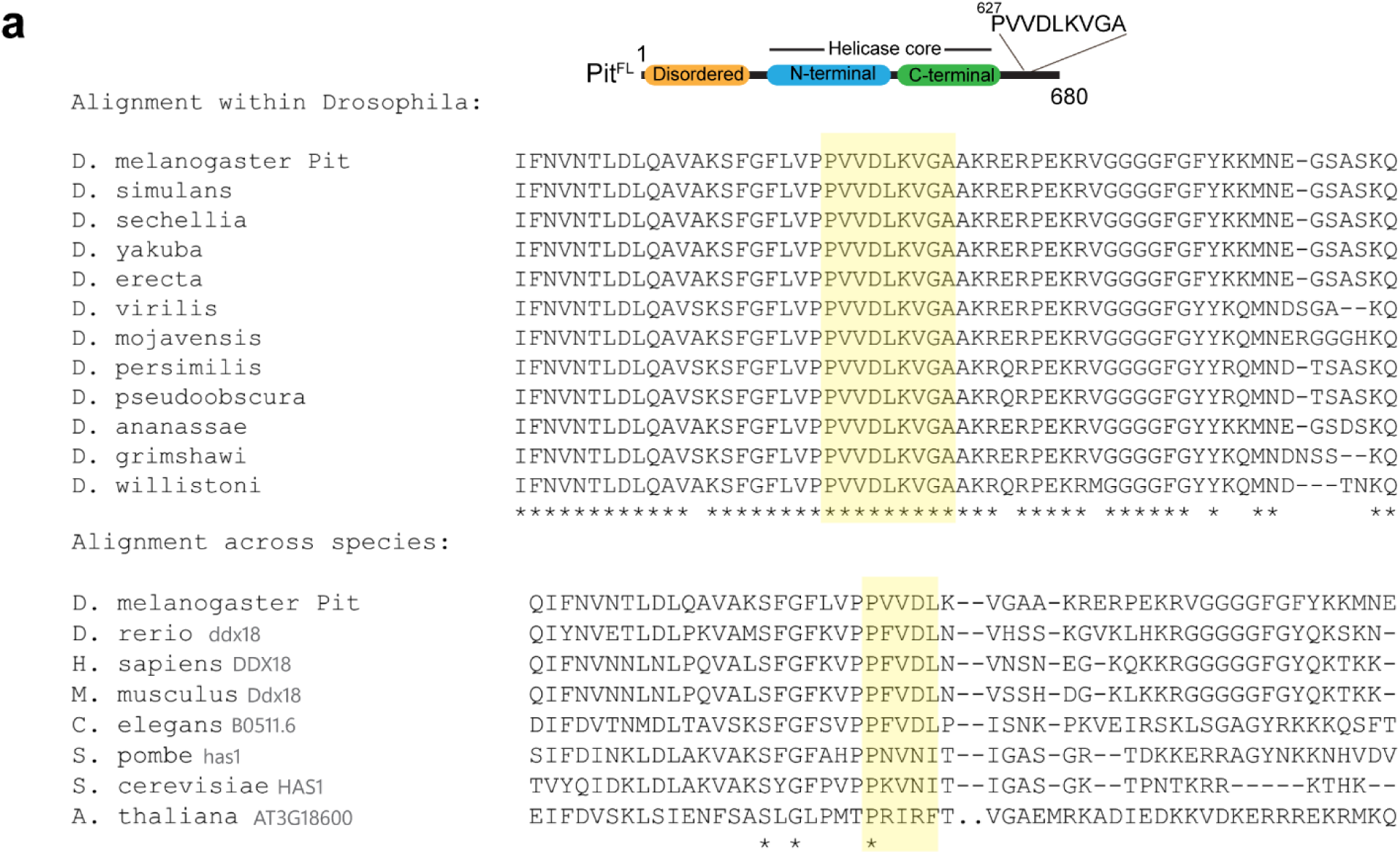
Pitchoune has a conserved PxVxL HP1a-interacting motif. (**a**) Evolutionary analysis to determine the conserved PxVxL motif in Pitchoune within *Drosophila* species (top) and across non-*Drosophila* eukaryotic model organisms (bottom).

**Extended Data Fig. 6:**
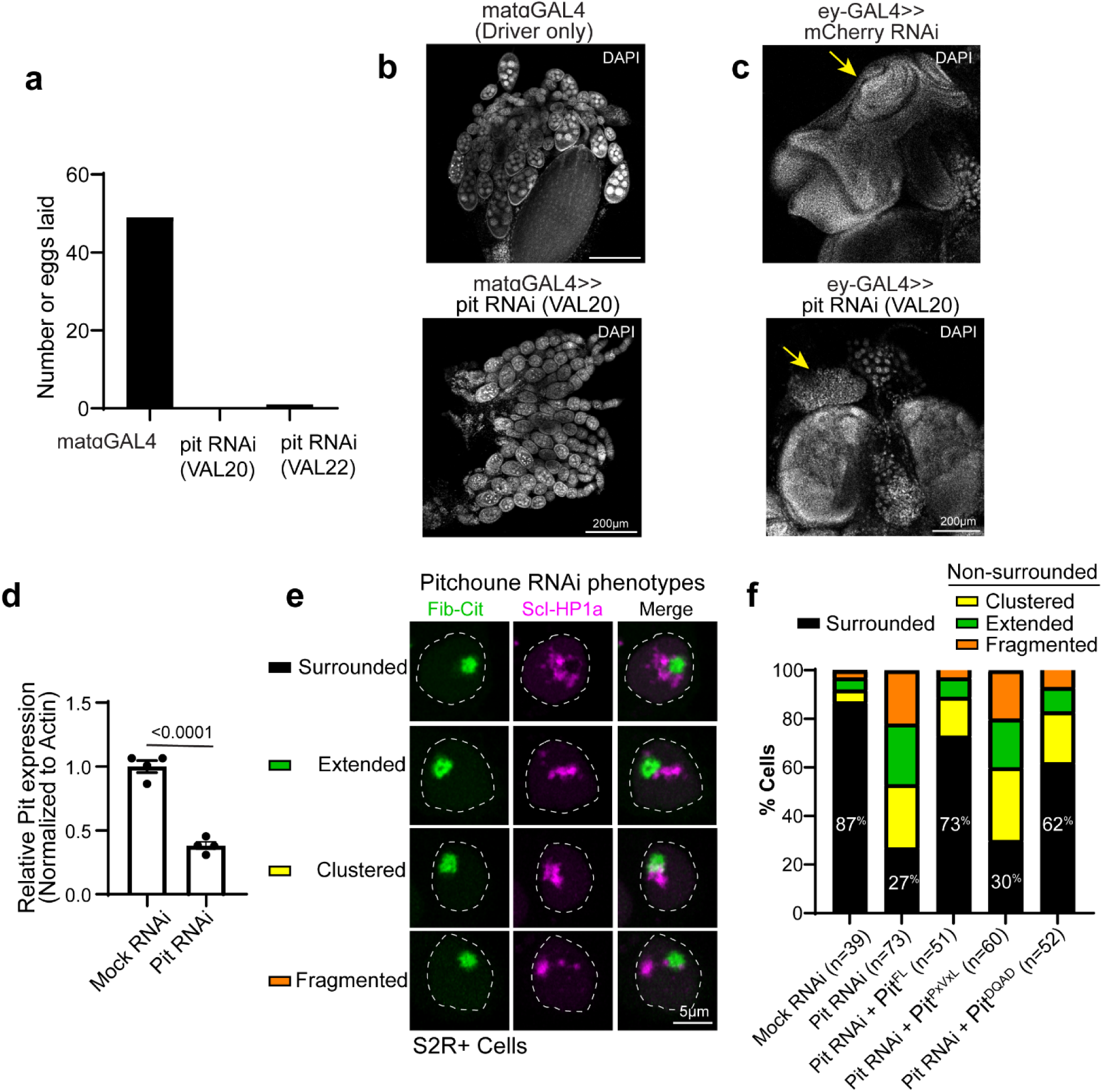
Developmental defects and HP1a disorganization phenotypes due to Pitchoune knockdown in Drosophila tissues and cultured cells. (**a**) Number of eggs laid on an apple juice plate after a 3hr collection in mataGAL4 (driver only) and mataGAL4 driving UAS-Pitchoune RNAi (VAL20 and VAL22 lines). (**b**) Representative images of dissected ovaries in control (mataGAL4, driver only) and after Pitchoune knockdown stained with DAPI. (**c**) Representative images of dissected eye antennal discs (yellow arrow) in control (ey-GAL4>mCherry RNAi) and after Pitchoune knockdown (ey-GAL4>pit RNAi, VAL20) stained with DAPI. (**d**) Quantitation of Pitchoune transcripts using qPCR to confirm the knock-down of Pitchoune, normalized to actin. Bar graphs depict mean ± s.e.m. n=4 biological replicates. (**e**) Representative nuclei transfected with Fib-Citrine and Scarlet-I-HP1a, showing four categories of HP1a distribution phenotypes observed after Pitchoune knockdown. (**f**) Quantification of the % nuclei with Surrounded, Extended, Clustered, and Fragmented HP1a phenotype after Pitchoune RNAi and its rescue with Pit^FL^, Pit^PxVxL^, and Pit^DQVD^.

